# Identification and Characterization of Novel *Faecalibacterium prausnitzii* Strains with Potential Pharmabiotic Applications

**DOI:** 10.64898/2025.12.30.697052

**Authors:** Olesya O. Galanova, Mikhail D. Akulinin, Daria A. Troshina, Alexey S. Kovtun, Maya V. Odorskaya, Vera V. Muravieva, Ksenia N. Zhigalova, Roman V. Izyumov, Alexey B. Gordeev, Bayr O. Bembeeva, Elena L. Isaeva, Alexey A. Vatlin, Tatiana V. Priputnevich, Valery N. Danilenko, Gennady T. Sukhikh

**Affiliations:** Laboratory of Bacterial Genetics, Vavilov Institute of General Genetics, Russian Academy of Sciences, Moscow, 119333, Russia; Moscow Center for Advanced Studies, 123592 Moscow, Russia; Faculty of Biotechnology, Lomonosov Moscow State University, Moscow, 119991, Russia; Microbiological Laboratory for Polymicrobial Infections and Biofilms, Kulakov National Medical Research Center for Obstetrics, Gynecology and Perinatology, Ministry of Health, Moscow, 117997, Russia; Institute of Environmental Engineering, RUDN University, 6 Miklukho-Maklaya St., 117198 Moscow, Russia

**Keywords:** Faecalibacterium prausnitzii, genome, pangenome, pharmabiotic potential, gut microbiome

## Abstract

*Faecalibacterium prausnitzii* is a dominant commensal bacterium in the human gut, widely known for its anti-inflammatory and immunomodulatory properties and currently considered a potential pharmabiotic for the treatment of various human diseases. In this study, ten new strains of *F. prausnitzii* were isolated from stool samples of Russian children and characterized using hybrid genome sequencing and metabolomic analyses. Comparative genomics revealed an open pangenome structure and significant strain-specific variability, reflecting the high adaptive potential of strains in humans and confirming the heterogeneity of this species previously described in the literature. Several isolates (fp1, fp5, fp7, fp9) showed increased butyrate production, while fp1 showed increased formate synthesis. *In silico* analysis confirmed the presence of genes associated with anti-inflammatory, antioxidant, and neuromodulatory metabolites, as well as the absence of antibiotic resistance genes. These results expand on previously described findings on the genetic diversity of *F. prausnitzii* and identify promising, safe strains with high pharmabiotics potential for future human therapy.

## 1. Introduction

*Faecalibacterium prausnitzii* is a Gram-positive, anaerobic bacterium and one of the most abundant commensals in the human colon, comprising up to 5% of the total bacterial population in healthy adults [1]. According to comparative genomic studies, the genomes of *F. prausnitzii* are characterized by high intraspecific variability, a low level of average nucleotide identity (approximately 85%) and significant plasticity [2,3].

The growing interest in *F. prausnitzii* is related to its important role in maintaining intestinal homeostasis. A marked decrease in the abundance of *F. prausnitzii* is considered an indicator of dysbiosis and is associated with a wide spectrum of diseases. Its depletion is consistently observed in inflammatory bowel diseases (IBD), including Crohn’s disease and ulcerative colitis, where its levels correlate inversely with disease severity [4]. Similarly, reduced levels have been reported in such conditions as colorectal cancer (CRC), irritable bowel syndrome (IBS), type 2 diabetes, obesity, and atopic dermatitis [5,6]. In metabolic disorders like obesity, this reduction contributes to decreased butyrate production and an elevated pro-inflammatory state [7]. Furthermore, dysbiosis characterized by a lack of *F. prausnitzii* has been linked to mental health disorders in animal models, potentially through its role in modulating the gut-brain axis [8]. Therefore, *F. prausnitzii* might offer a useful novel therapeutic approach to a variety of disorders.

*F. prausnitzii* is involved in the maintenance of gut homeostasis primarily through the production of short-chain fatty acids (ScFAs), such as formate, acetate, propionate and butyrate [9]. These ScFAs interact with receptors in various tissues to exert systemic signaling effects [10]. Butyrate and propionate also provide an epigenetic influence by inhibiting histone deacetylases and regulating gene expression [11]. Notably, butyrate serves as the primary energy source for colonocytes, enhances intestinal barrier function by upregulating tight junction proteins, and exerts potent immunomodulatory effects [12]. These anti-inflammatory properties are mediated through the inhibition of the NF-κB pathway and pro-inflammatory cytokines like IL-12 and IFN-γ, while stimulating the secretion of the anti-inflammatory cytokine IL-10 [13]. Beyond SсFAs, *F. prausnitzii* produces microbial anti-inflammatory molecules (MAM) and aromatic acids, such as shikimic and salicylic acids, which further contribute to dampening intestinal inflammation [14,15]. *F. prausnitzii* produces many of its beneficial metabolites by metabolizing dietary fibers. This activity is linked to a lower risk of microbial translocation and inflammation, which helps prevent gastrointestinal comorbidities [16]. Collectively, these multifaceted beneficial properties position *F. prausnitzii* as a leading candidate for the development of pharmabiotics—bacterial cells or their products with a proven pharmacological role in health and disease [17–19].

The practical application of *F. prausnitzii* in therapeutics is complicated by its high genetic and functional heterogeneity. Various strains exhibit significant variability in their metabolic capabilities, including the production of butyrate and other anti-inflammatory metabolites, as well as in their tolerance to oxygen exposure [20]. The composition and functional profile of an individual’s *F. prausnitzii* population are influenced by factors such as age, diet, and geography [21,22]. Importantly, some strains may harbor virulence factors or antibiotic resistance genes, necessitating rigorous safety screening when selecting probiotic candidates [23].

However, despite the well-documented anti-inflammatory properties of *F. prausnitzii*, its high genomic and metabolic heterogeneity remains a major limitation for therapeutic development [24]. Furthermore, population-specific variations—driven by age, diet, and geography—may significantly influence strain functionality [25]. To address these gaps, we isolated and characterized ten novel *F. prausnitzii* strains from the gut microbiota of healthy children in the Moscow region. Using a hybrid sequencing and metabolomic approach, we aimed to assess their genetic diversity, metabolic profiles, and pharmabiotic potential.

## 2. Materials and Methods

### 2.1. Biomaterial collection

Fecal samples were collected from children residing in the Moscow region of Russia using sterile containers. Samples were obtained immediately (within 1 minute) after natural defecation and placed into sterile collection containers. To maintain anaerobic conditions during transport, each container was placed into a zip-lock bag containing a gas-generating sachet (AnaeroGen Compact, Oxoid). The bags were hermetically sealed and transported to the laboratory for further processing.

### 2.2. Isolation and Cultivation of Faecalibacterium prausnitzii Strains

Fecal culturing was processed in an anaerobic chamber (BACTRON 300-2, Shel Lab, USA) under an atmosphere of a three-component gas mixture (N₂-80%; CO₂-10%; H₂-10%). Fecal samples were weighed and serially diluted ten-fold in phosphate-buffered saline (PBS) from 10⁻² to 10⁻⁹, and 0.1 ml aliquots of the 10⁻⁷ to 10⁻⁹ dilutions were plated onto Petri dishes with agarized nutrient media.

For this work we developed a solid culture medium for the cultivation of *F. prausnitzii* as it belongs to the group of obligate anaerobic microorganisms that are highly sensitive to oxygen. This nutrient medium for the isolation of strictly anaerobic, extremely oxygen-sensitive, short-chain fatty acid-producing microorganisms patented by Russian Federation Patent for Invention No. 2833071, issued: 14.01.2025; priority date: 24.05.2024.

Identification of the grown colonies was performed by MALDI-TOF MS using a Microflex mass spectrometer with MALDI BioTyper software version 4.1 (Bruker Daltonics, Germany). All microbial isolates that were not successfully identified by MALDI-TOF MS were identified by Sanger sequencing of the 16S rRNA gene fragment (V3-V4 regions). Pure cultures of *F. prausnitzii* were lyophilized and also preserved in liquid nutrient medium with the addition of a cryoprotectant (15% glycerol) at −80°C (Table 1).

**Table 1.** Description of biological sources and isolated strains.

| Strain N° | Age and gender of the child | Selection date | BioProject ID |
| --- | --- | --- | --- |
| fp1 | male child, 11 months of age | 29.03.2024 | PRJNA1120745 |
| fp2 | female child, 12 months of age | 11.04.2024 | PRJNA1120746 |
| fp3 | female child, 11 months of age | 16.04.2024 | PRJNA1116079 |
| fp4 | female child, 12 months of life | 15.04.2024 | PRJNA1116080 |
| fp5 | female child, 11 months of age | 26.04.2024 | PRJNA1116081 |
| fp6 | female child, 11 months of age | 26.04.2024 | PRJNA1120742 |
| fp7 | male child, 11 months of age | 27.05.2024 | PRJNA1122146 |
| fp8 | female child, 12 months of age | 22.05.2024 | PRJNA1122147 |
| fp9 | male child, 11 months of age | 31.05.2024 | PRJNA1122154 |
| fp10 | female child, 10 months of age | 03.06.2024 | PRJNA1122155 |

### 2.3. Genomic DNA Extraction and Sequencing

Genomic DNA was extracted using the QIAamp DNA Stool Mini Kit, following the manufacturer’s protocol. DNA concentration and purity were assessed with a NanoDrop 2000 spectrophotometer. High-quality DNA samples were underwent using Illumina HiSeq 2500 for short reads and Oxford Nanopore MinION for long reads. Sequencing libraries were prepared according to the respective protocols.

### 2.4. Genome Assembly

Quality assessment of the raw sequencing data was performed using FastQC [26] Adapter trimming and read quality enhancement were carried out with Trimmomatic (v.0.39) [27] for short reads and Porechop (v.0.2.4) for long reads [28]. *De novo* genome assembly was conducted using long reads via the Flye assembler (v.2.9.5) [29]. To polish the initial assembly, short reads were mapped to the draft genome using BWA (v.0.7.18) and processed with SAMtools (v.1.21) [30]. The resulting assembly and alignment (BAM) files were then used as input for Pilon (v.1.2.4) to correct errors and fill gaps [31]. The quality of the final assembly was assessed using QUAST (v.5.3.0) [32] and BUSCO (v.1.0.0) [33]. Genome annotations were performed using Prokka (v.1.14.5) [34].

### 2.5. Taxonomic annotation of the assembled strains

#### 2.5.1. Nucleotide identity

To confirm the species identity of the ten studied strains, a comprehensive nucleotide-level comparative genomic analysis was performed using multiple genome similarity metrics, including average nucleotide identity (ANI), ANIm, ANIb, and tetranucleotide frequency correlation (TETRA).

ANI analysis was conducted using FastANI (v.1.34) to compare the studied genomes with reference *Faecalibacterium prausnitzii* genomes retrieved from the NCBI database [35]. The analysis was performed using default parameters, ≥ and an ANI threshold of 95% was applied as the criterion for assignment to the same species.

The ANIm analysis was performed using the pyani package (v.0.3.2) using MUMmer-based alignment [36]. This analysis was performed only on the complete genomes. ANIm uses anchor sequence alignment to calculate nucleotide similarity.

The ANIb analysis was performed using the pyani package and BLAST+ algorithm for pairwise comparison of genomes [37]. Like ANIm, this analysis was performed only on the complete genomes. ANIb uses the pairwise alignment of BLAST to calculate the percentage of identity of conserved regions of genomes.

In addition, TETRA analysis was performed using the pyani package to calculate the correlation coefficients of tetranucleotide frequencies. This method provides confirmation of taxonomic affiliation based on the characteristics of the use of nucleotides in genomes.

The threshold for all nucleotide metrics to confirm membership in the *F.prausnitzii* species was 0.9 or 90%

#### 2.5.2. 16S rRNA species identification

For species identification using 16S rRNA sequences, a reference database was constructed. Full-length 16S rRNA gene sequences of *Faecalibacterium prausnitzii* were downloaded from the NCBI database via Entrez-Direct. The initial dataset was curated by applying a length filter (1,200 to 1,700 bp), followed by dereplication and *de novo* chimera removal using VSEARCH (v.2.30.0) [38]. The alignment of the novel genomes to the 16S rRNA *Faecalibacterium prausnitzii* sequence base was performed. High-confidence alignment hits were identified by applying filtering criteria: an e-value cutoff of 1e-20, a minimum query coverage of 85%, and a minimum sequence identity of 85%.

### 2.6. Nucleotide sequence diversity among isolate genomes

#### 2.6.1. Diversity among novel strains

To identify homologous genomic regions between strain assemblies, FastANI (v1.34) was used. A minimum ANI threshold of 80% was applied for inclusion; genomic regions with ANI values below this threshold were excluded from subsequent analyses. The results of the pairwise genomic comparisons were visualized using Circos [39].

#### 2.6.2. Pangenome analysis

For pangenome analysis, a total of 27 *F. prausnitzii* strains were included in this study: 17 complete genomes retrieved from the National Center for Biotechnology Information (NCBI) database (**Table 2**) and 10 strains obtained in this study (**Table 1**). All genomes were annotated using Prokka. Pangenome construction and analysis were performed using Panaroo (v.1.5.2) [40] run in “--clean-mode” strictly to minimize annotation errors. Homologous genes were clustered using MMseqs2 with a threshold of 80% amino acid sequence identity [41]. Core gene alignments were generated using MAFFT [42]. Pan- and core-genome curves were visualized using PanGP (v.1.0.1) [43] with the gene presence/absence matrix as input. Functional annotation of the gene clusters was performed using eggNOG-mapper (v.5.0.2) [44] against the Clusters of Orthologous Groups (COG) database [45]. Visualization of COG gene characterization was generated using the ggplot2 package in R.

**Table 2.**
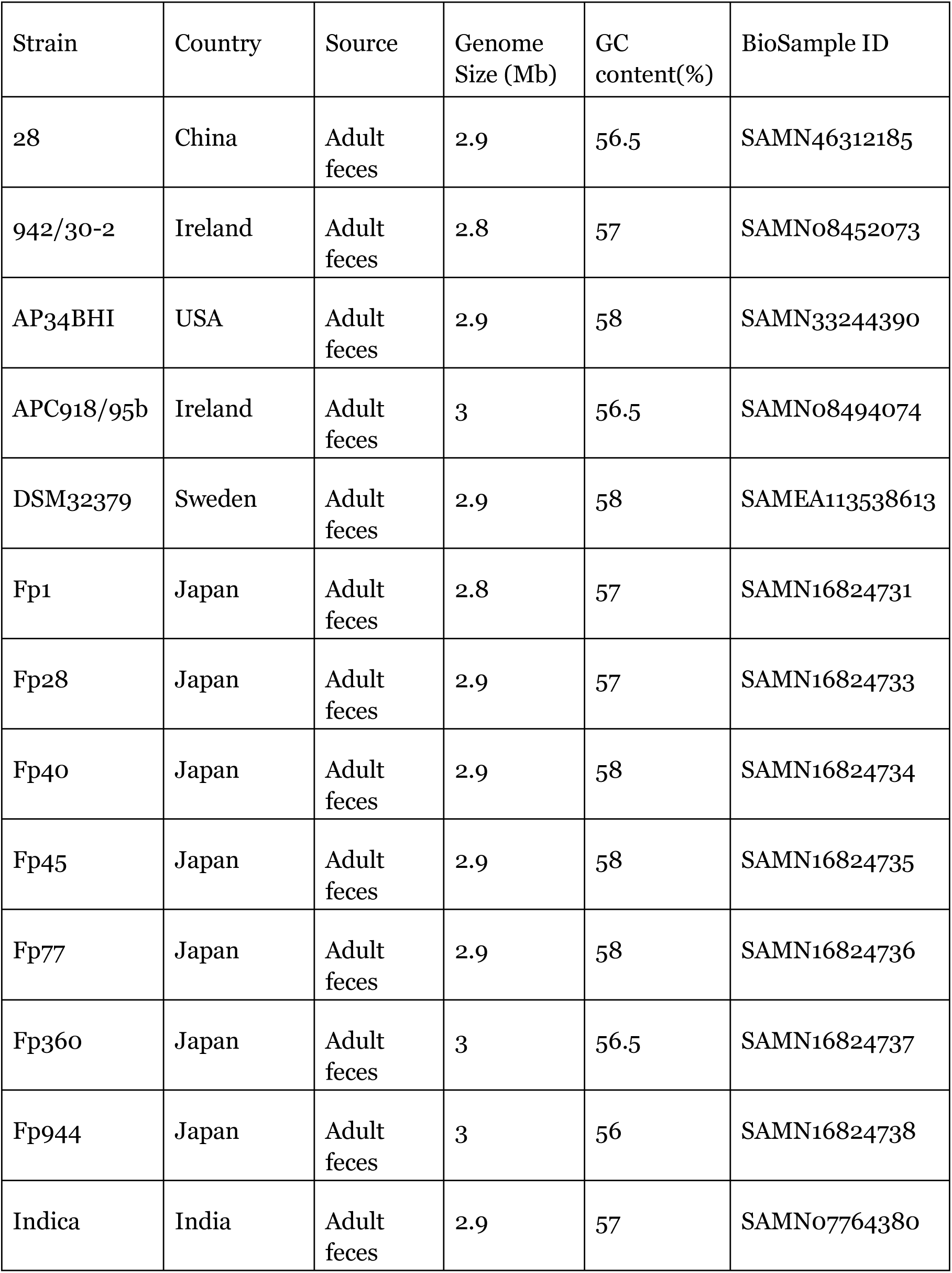
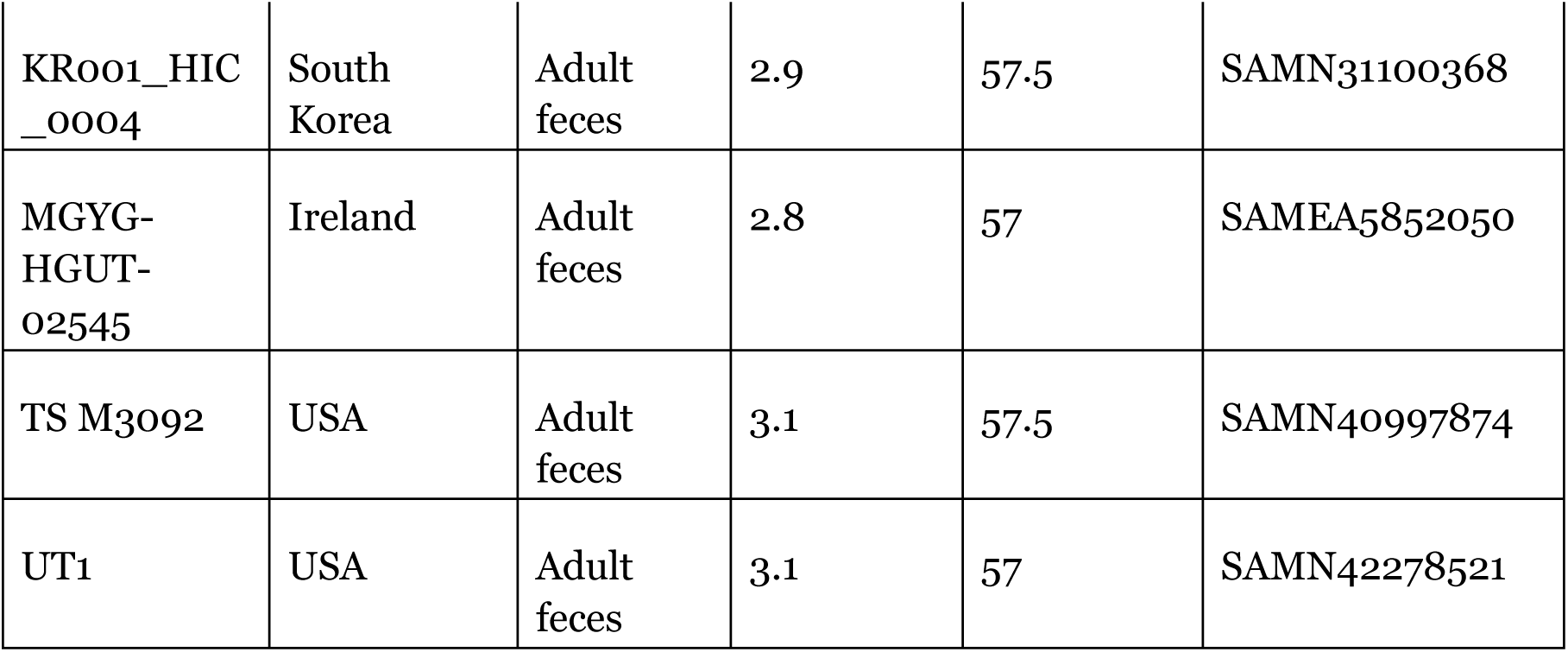
General properties of *F. prausnitzii* strains with available complete genome sequences.

#### 2.6.3. Phylogenetic analysis

Maximum-likelihood phylogenetic trees were constructed to assess the clustering of the newly isolated strains relative to reference *Faecalibacterium prausnitzii* strains retrieved from the NCBI database. A maximum-likelihood phylogenetic tree was inferred using IQ-TREE (v.2.2.0) based on a concatenated core-gene alignment [46].

The best-fitting nucleotide substitution model was selected by the ModelFinder option “-m MFP”. Node support was assessed using 1,000 bootstrap replicates “-bb 1000” and 1,000 SH-aLRT replicates “-alrt 1000”. The aligned core genes also were processed using snp-sites (v.2.5.1) to identify SNPs [47]. Subsequently, the best-fit substitution model was selected based on the BIC criterion using modeltest-ng (v.0.1.7), resulting in GTR+G4 as the optimal model [48]. Phylogenetic tree reconstruction was performed using the maximum likelihood method implemented in raxml-ng (v.1.2.2) with the GTR+FO+G4m model and 1000 bootstrap replicates [49]. The phylogenetic trees were visualized in iTOL [50].

#### 2.6.4. Reference genes catalog development

The catalog of amino acid sequences implicated in the metabolism of compounds possessing antioxidant, neuromodulatory, and immunomodulatory activities was developed and published earlier [51]. This catalog was used as the foundation for characterizing the pharmabiotic potential of *F.prausnitzii* strains from our collection. The list of target metabolites was. For each compound, corresponding KEGG identifiers were retrieved [52], and enzyme commission (EC) numbers associated with their biosynthesis or utilization were identified. Amino acid sequences of the corresponding enzymes were obtained from the UniProt database and filtered to include representatives from the 46 most prevalent bacterial genera of the human gut microbiome [53]. Newly identified homologs were subsequently incorporated into the previously established catalog of gene orthologs.

#### 2.6.5. Analysis of homologs

To characterize our novel *F. prausnitzii* strains, compare their annotated amino acid sequences against a reference sequence catalog using BlastP. Matches were retained only if they exhibited >60% sequence identity and a length difference of <10% compared to the reference sequences. Homology searches for drug resistance and virulence genes were conducted using the same criteria. Additionally, we performed the comparison of novel *F. prausnitzii* strains with each other and *F. prausnitzii* strains with complete genome sequences available in NCBI. The analysis was performed using Blast+ and homologs sequences from the previous step.

### 2.7. Metabolomic short-chain fatty acid analysis

#### 2.7.1. Sample preparation

1. *F. prausnitzii* strains isolated from fecal samples were initially cultured on solid nutrient medium for 48 h. The resulting pure cultures were then grown in a liquid nutrient medium for metabolomic profiling. The liquid medium used was LYHBHI [54] based on Brain Heart Infusion broth (BHI, HiMedia, India) (37 g/L), supplemented with yeast extract (FBUN SRC PMB, Russia) (5 g/L), maltose (FBUN SRC PMB, Russia) (1 g/L), cellobiose (CDH Central Drug House, India) (1 g/L), and cysteine (AppliChem, Germany) (0.5 g/L).
2. *F. prausnitzii* initial inoculum was prepared to an optical density of 2 McFarland standard and cultured under anaerobic conditions at 37 °C in an anaerobic chamber (N₂-80%; CO₂-10%; H₂-10%). Changes in the optical density of the inoculum were measured at three time points: 0 hours, 6 hours (late exponential growth phase), and 24 hours (stationary growth phase) to assess culture growth kinetics. At each time point, a series of ten-fold dilutions of the inoculum were prepared in the same medium, and aliquots from the 10⁻⁵ to 10⁻⁷ dilutions were plated onto a solid nutrient medium to determine the number of viable cells (Colony Forming Units - CFU) [54,55]. The metabolomic profile of culture supernatants was analyzed after 24 hours of cultivation (during the stationary growth phase).

#### 2.7.2. Samples of culture media

An aliquot of 100 µL of the culture medium was transferred into a 500 µL plastic microcentrifuge tube. 10 µl of a combined solution of internal standards and 5 µl of hydrochloric acid solution (2 M) were added to the sample. After brief mixing, 200 μL of methyl tert-butyl ether (MTBE) was added to the mixture. Liquid-liquid extraction was then performed for 20 minutes under vigorous agitation. Phase separation was achieved by centrifugation at 15,000 rpm for 5 minutes. A 100 μL aliquot of the organic phase (upper layer) was transferred into an autosampler vial equipped with a glass insert for subsequent analysis.

#### 2.7.3. Reagents and standards

For the analysis of short-chain fatty acids (ScFAs) concentration in culture media, acetic acid-d4 (AA-d4) was used as an internal standard for the quantification of acetic and formic acids, and butyric acid labeled with two carbon-13 atoms (13C2-BA) was used for the quantitative analysis of the remaining acids (propionic, butyric, isobutyric, valeric, isovaleric, 2-methylbutyric, 3-methylbutyric, caproic, and caprylic acids).

MTBE of high purity (HPLC-grade) was used for the extraction. Acetonitrile (ACN) served as the solvent for preparing the internal standard solutions. A hydrochloric acid solution (2 M) was used for sample acidification. All solutions were prepared using deionized water (Milli-Q grade).

The ISTD1 solution was prepared by dissolving 5.7 μL of AA-d4 in 10 mL of acetonitrile. The ISTD2 solution was prepared by dissolving 100 mg of 13C2-BA in 10 mL of acetonitrile. A pooled internal standard (ISTD) solution was formed by mixing 991 μL of ISTD1 and 9 μL of ISTD2, providing final approximate internal standard concentrations of 10^4^ μM and 10^3^ μM, respectively.

#### 2.7.4. Calibration solutions

Calibration mixtures were prepared in a water base. To each 100 μL aliquot of the standard compound mixtures, 10 μL of the internal standard solution and 5 μL of hydrochloric acid (2 M) were added, followed by liquid-liquid extraction using 200 μL of MTBE.

The calibration levels were as follows: 1000, 5000, 10000, and 50000 μM for acetic and formic acids; 500, 2500, 5000, and 25000 μM for propionic acid; 100, 500, 1000, and 5000 μM for butyric and isobutyric acids; and 100, 500, 1000, and 5000 μM for valeric, isovaleric, 2-methylbutyric, 3-methylbutyric, caproic, and caprylic acids.

#### 2.7.5. Gas chromatography

Chromatographic separation was performed using an Agilent 7890B gas chromatograph equipped with an HP-FFAP column (25 m × 0.32 mm, 0.5 µm film thickness). The injector and GC-MS transfer line temperatures were maintained at 250 °C. The injection volume was 1 µL. The analysis was conducted in split mode with a ratio of 10:1. The carrier gas was helium with a purity of at least 99.9999%, at a constant flow rate of 1.7 mL/min. The oven temperature program was as follows: initial isothermal hold at 60 °C for 1.5 min; ramping at a rate of 20 °C/min to 240 °C; followed by a final hold at 240 °C for 3 min.

#### 2.7.6. Mass spectrometric detection

Detection was performed using an Agilent 5977B mass spectrometer with electron impact ionization (70 eV). The ion source temperature was set at 230 °C, the quadrupole at 150°C, and the transfer line at 250 °C. Detector activation was delayed for 4 minutes after the start of the chromatographic analysis to prevent damage to the ion source from solvent exposure. Ion detection was conducted in Selected Ion Monitoring (SIM) mode. For quantitative analysis, the following ions were used: m/z 60.0 for acetic, butyric, valeric, isovaleric, 3-methylvaleric, caproic, and caprylic acids; m/z 74.0 for propionic, 2-methylvaleric, and 2-methylbutyric acids; m/z 63.0 for AA-d4; m/z 62.0 for 13C2-BA.

#### 2.7.7. Data processing

Chromatographic data processing was performed using MassHunter Quantitative Analysis software. Calibration curves were constructed based on the ratio of the compound peak area to the corresponding internal standard peak area. Linear approximation was achieved by the least squares method. The coefficient of determination (R²) exceeded 0.99. The limit of detection (LOD) was calculated using the formula LOD = 3.3 × (σ / b), where σ is the standard deviation of the y-intercept and b is the slope of the calibration curve. The limit of quantification (LOQ) was calculated as LOQ = 3 × LOD. Chromatograms of standard solutions were used to confirm compound identification and verify the sensitivity of the method.

## 3. Results

### 3.1. Genomes assembly

Ten newly isolated strains of *Faecalibacterium prausnitzii* obtained from children in the Moscow region were assembled and characterized to assess their genomic diversity and assembly quality.

Assembly of the sequenced reads resulted in four complete genomes and six high-quality draft genomes. The average genome length and GC-content were 3.0 Mb and 56%, respectively (Table 3).

**Table 3.** General properties of novel *F. prausnitzii* strains genomes.

| Strain | Genome size (Mb) | Contigs number | GC content (%) | BioSample ID, NCBI |
| --- | --- | --- | --- | --- |
| fp1 | 3,1 | 1 | 56,9 | SAMN45776701 |
| fp2 | 3,1 | 1 | 56,2 | SAMN46972959 |
| fp3 | 3,0 | 5 | 56,1 | SAMN46972960 |
| fp4 | 2,9 | 3 | 57,2 | SAMN46972986 |
| fp5 | 3,0 | 1 | 56,2 | SAMN46973203 |
| fp6 | 2,6 | 4 | 56,1 | SAMN46973473 |
| fp7 | 3,1 | 1 | 56,5 | SAMN45804049 |
| fp8 | 2,8 | 6 | 55,7 | SAMN45804228 |
| fp9 | 3,0 | 4 | 56,6 | SAMN45804231 |
| fp10 | 3,0 | 4 | 56,6 | SAMN45804237 |

### 3.2. Taxonomic annotation of the assembled strains

#### 3.2.1. 16S rRNA identification

A total of 162 16S rRNA sequences of *F. prausnitzii* were downloaded from NCBI. Following the removal of duplicate and chimeric sequences, 117 sequences were retained for further analysis. Comparative analysis of the sequences of the 16S rRNA gene with the collected genomes revealed at least one corresponding sequence of the 16S rRNA gene of F. prausnitzii in each strain with sequence identity ≥95%. The resulting 16S rRNA gene sequences are presented in Figure 1.

**Figure 1.**
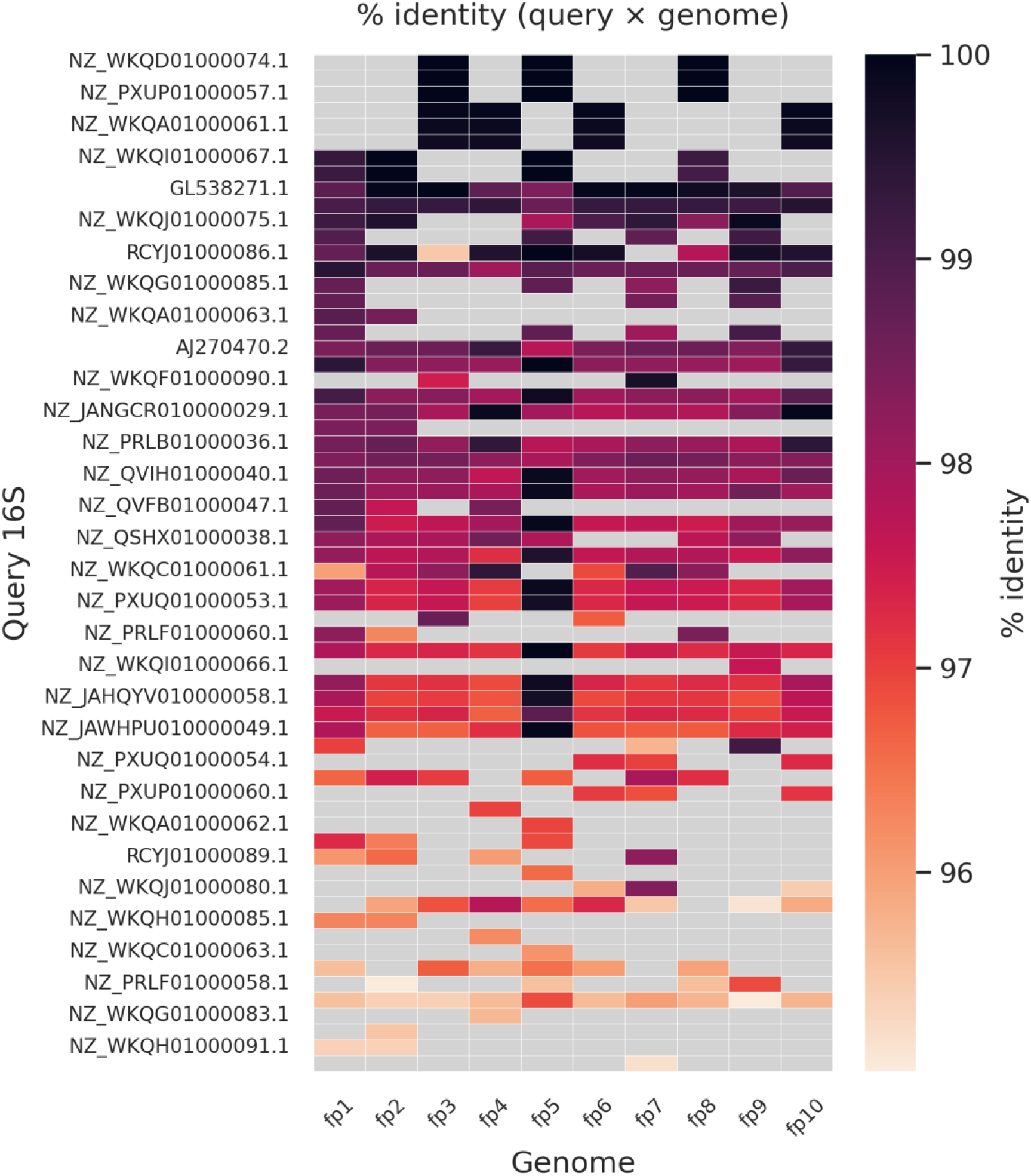
The heatmap illustrates sequence identity between the 16S rRNA sequences from NCBI and the assembled *F.prausnitzii* genomes. The X axis represents genomes, the Y axis represents 16S rRNA sequences from the NCBI database. The color scale represents the percent identity (95-100%). Gray squares mark empty numbers.

#### 3.2.2. Average nucleotide identity of assembled novel strains to NCBI strains *F.prausnitzii*

The analysis was carried out using the genomes of *F.prausnitzii* strains obtained from the NCBI database. Several genome comparison methods based on the similarity of nucleotide sequences were used (fastANI, ANIm, ANIb, TETRA). All newly assembled genomes were consistently classified as *F. prausnitzii* by each of the applied methods. For ANIm and ANIb only complete reference genomes were used.

Pairwise comparisons between our isolates and representative *F. prausnitzii* genomes from NCBI yielded high similarity values across multiple metrics (FastANI, ANIm/ANIb ≈0.97–1.00, tetranucleotide correlation 0.91–0.99). All these values are above the threshold value of 0.9. However, the top hit reference genome varied depending on the metric used. This discrepancy might reflect methodological differences between ANI implementations and heterogeneity in the reference genomes, rather than true incompatibility between metrics (Table 4).

**Table 4.** Nucleotide identity metrics of the comparison of the novel genomes with the genomes of novel genomes and genomes of *F.prausnitzii* strains from NCBI.

| Strain | FastANI | ANIm | ANiB | TETRA |
| --- | --- | --- | --- | --- |
| fp1 | 0.97 | 0.97 | 0.97 | 0.98 |
| fp2 | 0.97 | 1.0 | 1.0 | 0.96 |
| fp3 | 0.99 | 0.99 | 0.99 | 0.93 |
| fp4 | 0.99 | 1.0 | 1.0 | 0.99 |
| fp5 | 0.99 | 1.0 | 1.0 | 0.91 |
| fp6 | 0.99 | 1.0 | 1.0 | 0.91 |
| fp7 | 0.97 | 1.0 | 1.0 | 0.93 |
| fp8 | 0.97 | 1.0 | 1.0 | 0.91 |
| fp9 | 0.97 | 0.97 | 0.97 | 0.93 |
| fp10 | 0.98 | 1.0 | 1.0 | 0.99 |

### 3.3. Nucleotide sequence diversity

#### 3.3.1. Comparison of the assembled genomic sequences

Next we performed pairwise comparisons among the novel strains themselves to assess their intra-species genomic relatedness. Based on the comparison of the obtained genomes using the ANI metric, the strains can be divided into three groups: highly similar (above 95%), moderately similar (90% to 95%), and with low similarity (80% to 90%).

The following genome pairs were classified as highly similar: fp1/fp2, fp4/fp10, fp3/fp7, fp3/fp9, fp7/fp9, fp6/fp8. The following strain pairs were moderately similar: fp6/fp7, fp6/fp9, fp3/fp6, fp2/fp6, fp8/fp9, fp3/fp8, fp7/fp8. For the remaining strain pairs, the ANI value was below 90%, indicating low similarity.

The circos plots shown on Figure 2 represent the genomes’ homology by color bands. The “white” zones on the graphs indicate that the homology between the strains is less than 80%.

**Figure 2.**
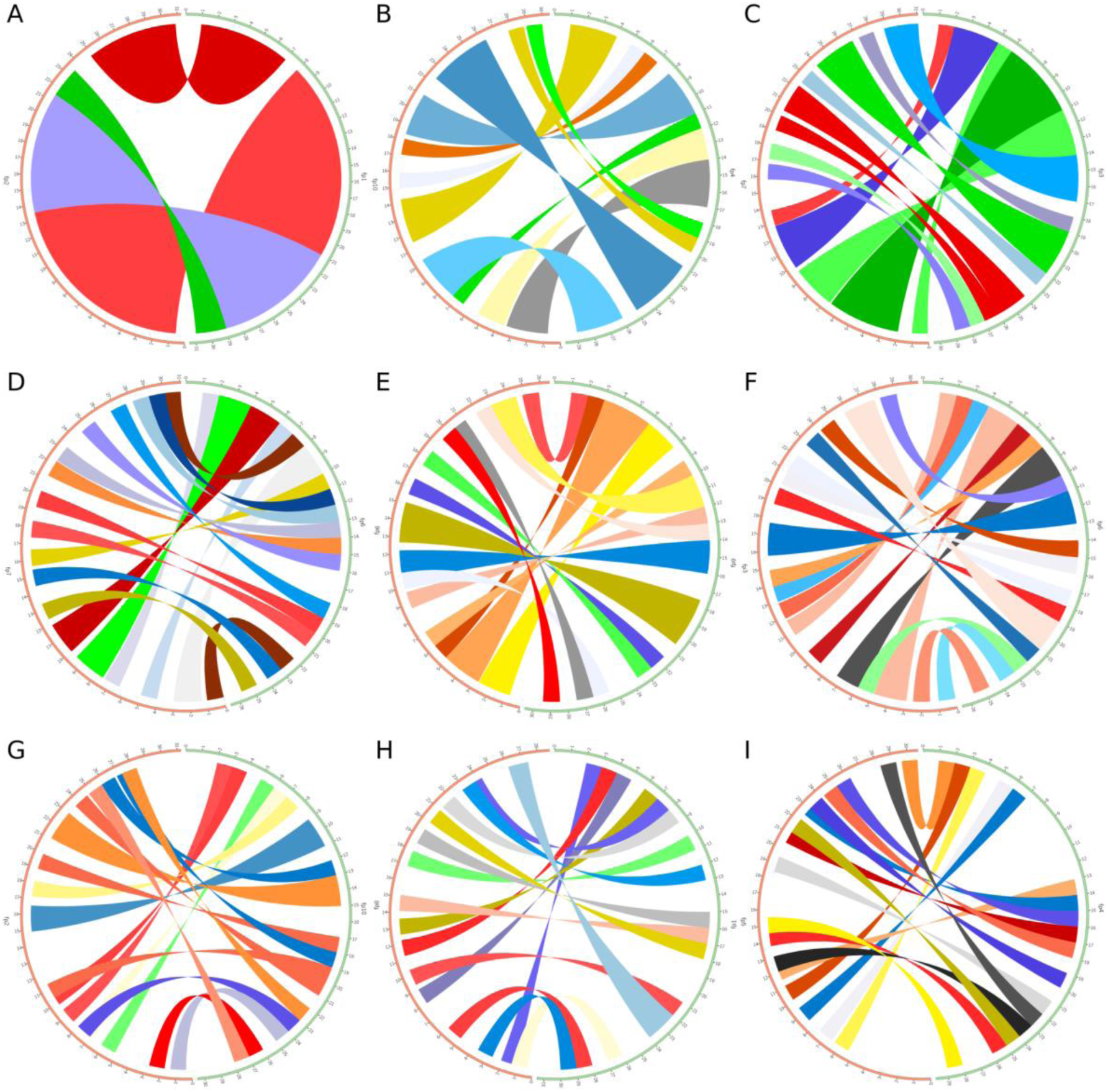
Circos plots representing the similarity of strains by pairs. Right and left halves locate a genome strain. A. Compare fp1 and fp2; B. Compare fp4 and fp10; C. Compare fp3 and fp7; D. Compare fp6 and fp7; E. Compare fp6 and fp9; F. Compare fp3 and fp6; G. Compare fp2 and fp10; H. Compare fp1 and fp8; I. Compare fp4 and fp5.

As we can see fp1 has the greatest homology with fp2 among all other compare pairs; its ANI metric is 97%. fp4/fp10 and fp3/fp7 have ANI about 96,5%. fp6/fp7, fp6/fp9, fp3/fp6 have ANI about 91%; in picture 2 (D-F) this is evident by the thinner color bands and more white gaps. For comparison pairs fp2/fp10, fp1/fp8, fp4/fp5, the ANI metric is about 82%; the figures show that Fig. 2 (G-I) contains quite a lot of white areas, which indicate regions with ANI below 80%. At the same time, the lengths of the genomes differ slightly.

#### 3.3.2. Pangenome analysis reveals high genetic diversity in Faecalibacterium prausnitzii

Combined analysis of 27 *F. prausnitzii* genomes revealed an open pangenome consisting of 12,784 gene clusters. The core genome represented the genes that are present in every single genome and contained 596 gene clusters (5% of pangenome size). The accessory genome denoted the set of genes that included some, but not all, genomes of the species and consisted of 8442 gene clusters (66% of pangenome size). The unique genome defines the set of genes that are found in only one genome (strain-specific gene) 3746 gene clusters (29% of pangenome size). The distribution of gene clusters in each strain is shown in figure 1. The number of accessory genes varied from 1253 to 1871, and for the unique ones - from 0 to 397. Some strains from the pangenome (Fp40, Fp45, 942/30-2 and MGYG-HGUT-02545) did not have any unique genes (Fig 3.). The absence of unique genes usually indicates that the strain is highly similar to the other genomes in the dataset.

**Figure 3.**
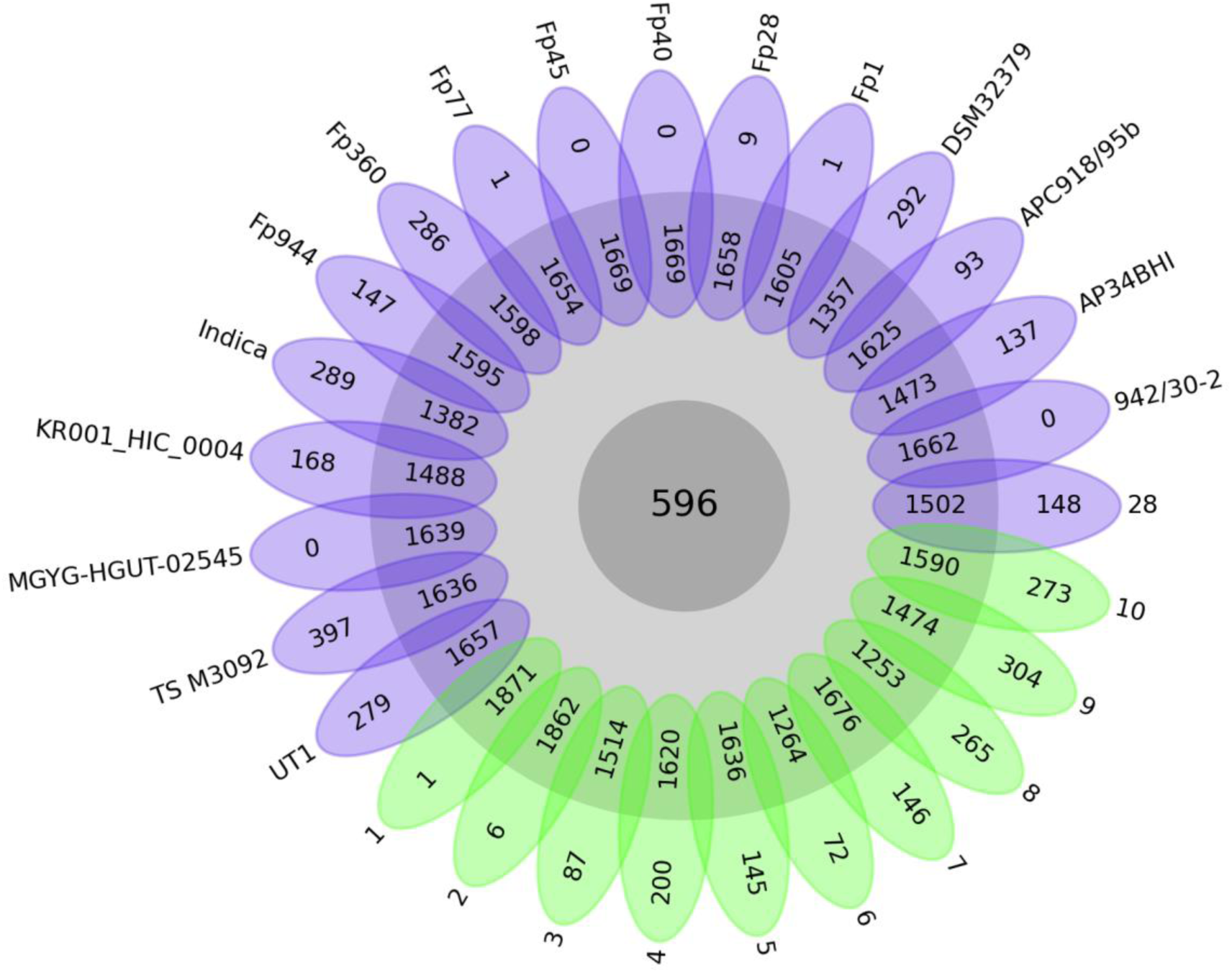
The flower plot illustrates the structure of the pangenome, with the core genome at the center, accessory genes in intermediate regions, and unique genes forming the petals. Strains are color-coded by source (blue: strains from NCBI; green: novel 10 strains).

To understand the relationships between pan- and core-genome size and the strain numbers of *F. prausnitzii* we plotted the fitted curves of the pan-genome profiles for 27 strains. The pangenome size curve did not reach the horizontal asymptote. This is the primary characteristic of an open pangenome [56]. This indicates that sequencing more genomes from this species will lead to the discovery of more new genes. Core-genome size curve is decreasing but appears to be approaching a stable plateau. This suggests that after analyzing a certain number of genomes (approximately 15-20 in this case), the essential, shared core genome has been reliably identified. Adding more genomes only trim a few more variable genes from the core set (Fig.4 A).

**Figure 4.**
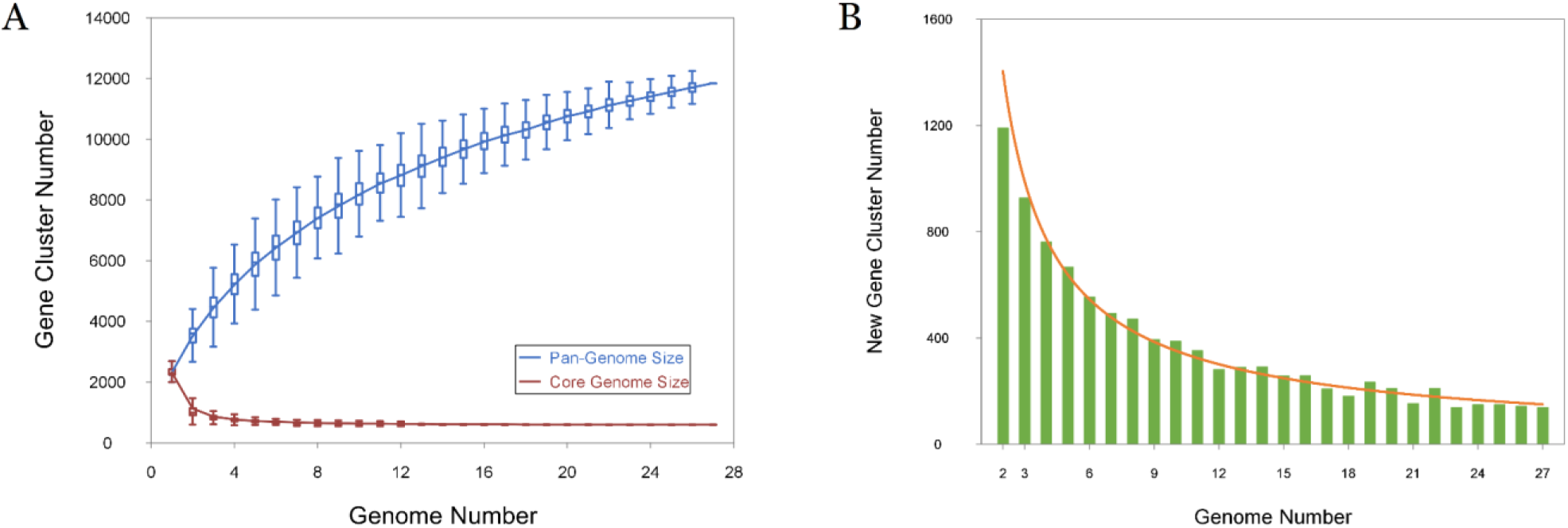
**A.** The pangenome curves of 27 *F. prausnitzii* strains. Fitting for the pan-genome profile curve (blue) is y=7913.03x^0,25^-5782.56 (R^2^=0,999107). Fitting for the core-genome profile curve (red) is y=4856.82e^-1,05x^+623.53 (R^2^=0,989176). **B.** The new gene cluster number plot for 27 *F. prausnitzii* strains. Fitting for New Gene profile curve (orange) y=2548.23x^-0,86^ (R^2^ = 0,971158).

The plot indicating the number of new gene clusters was also constructed. It shows a clear downward trend (Fig.4 B). The first two genomes added contribute a very high number of new gene clusters (over 1200 for the 2nd genome). However, by the time the 27th genome is added, it contributes very few new genes (likely well under 100). This indicates that the majority of the *F. prausnitzii* unique gene families have been discovered.

The pangenome analysis reveals a species with a stable core of essential genes and a large, open, and diverse accessory genome, which is a common signature for adaptable bacterial species like *F.prausnitzii*. To gain insight into the functional features of the pan-genome, we characterized genes using the Clusters of Orthologous Groups (COG) database (Fig. 5).

**Figure 5.**
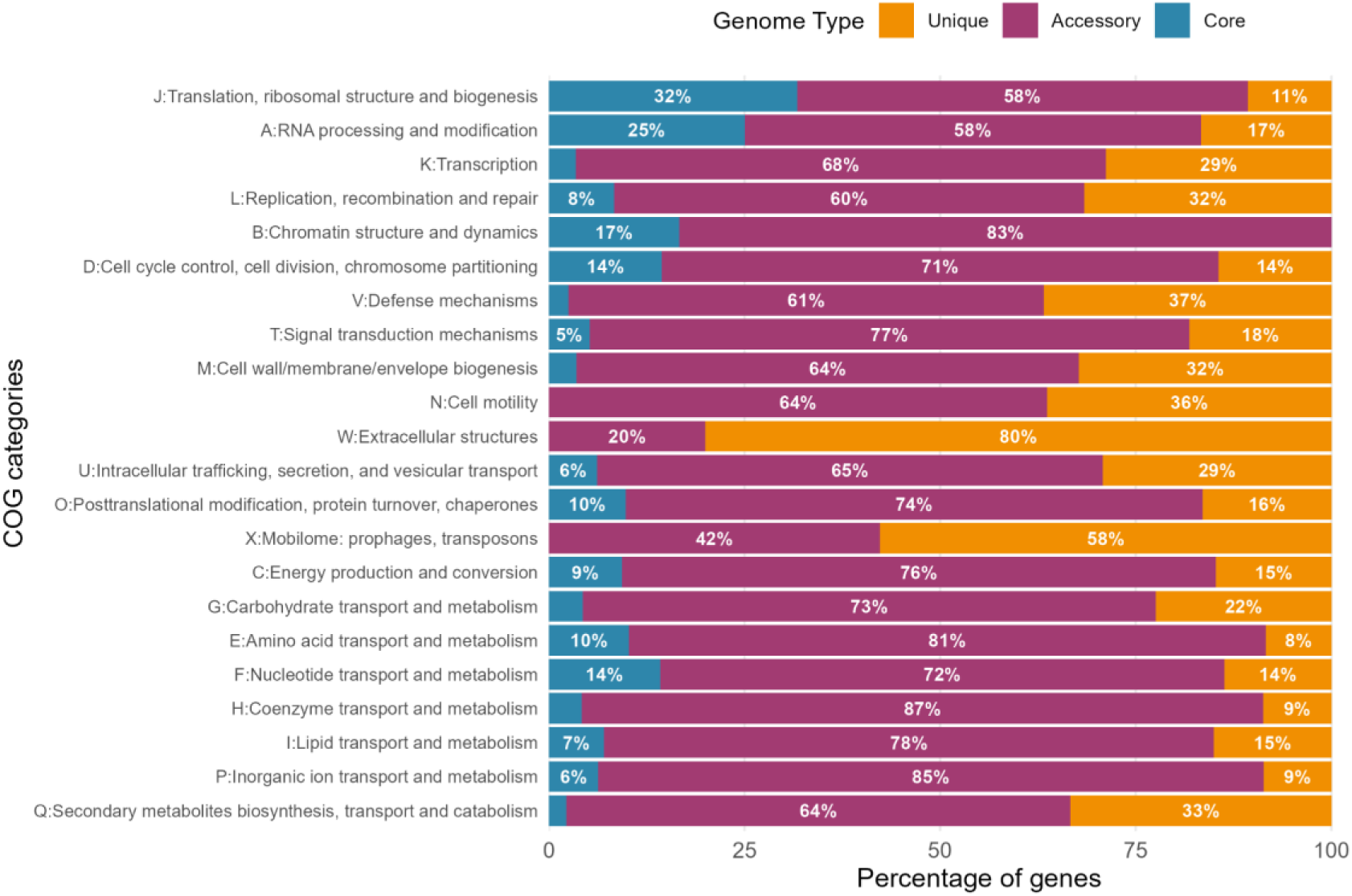
The proportion of genes of pan-genome based on COG categories.

A high proportion (43.3%) of the pan-genome comprised genes encoding proteins with unknown functions or no homologs outside the genus (9.7% of all core genomes, 40.7% - of all accessory genomes, 54.6% - of all unique genomes). The core genome was significantly enriched in essential housekeeping functions, including translation, ribosomal structure and biogenesis (J, 32%), DNA replication, recombination and repair (L, 9%), and cell cycle control (D, 14%). Conversely, the accessory genome was predominantly composed of genes related to environmental adaptation and niche specialization. Major functional categories included carbohydrate transport and metabolism (G, 73%), signal transduction mechanisms (T, 77%), and inorganic ion transport (P, 85%). This pattern reflects the evolutionary adaptation of *F. prausnitzii* to diverse gut environments and dietary substrates. The unique genome was characterized by a high abundance of mobile genetic elements (mobilome: prophages, transposons, X, 58%), genes encoding defense mechanisms (V, 37%) and genes encoding extracellular structures (W, 80%) indicating recent horizontal gene transfer events and strain-specific evolutionary adaptations.

#### 3.3.3. Phylogenetic analysis

The maximum-likelihood phylogenetic tree constructed from the concatenated core-gene alignment showed that the studied *F. prausnitzii* strains were successfully clustered with reference strains from the RefSeq database (Figure 6).

**Figure 6.**
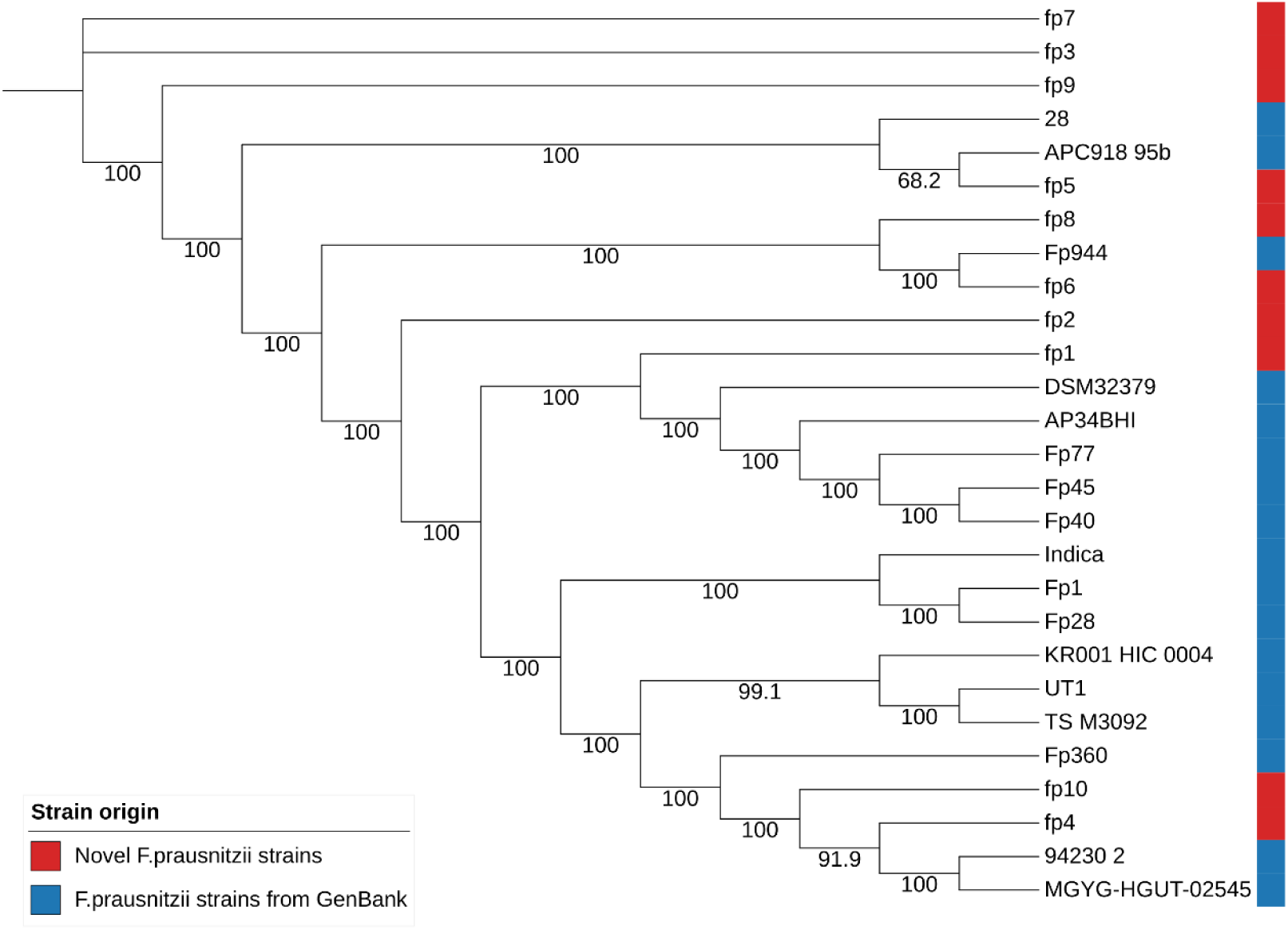
Maximum-likelihood phylogenetic tree based on the concatenated core-gene alignment of *F. prausnitzii* genomes. Colors indicate the origin of each strain, and numbers correspond to branch support values (bootstrap).

The novel strains fp4 and fp10 were located in the same cluster with *F. prausnitzii* 94230_2, *F. prausnitzii* MGYG-HGUT-02545, and *F. prausnitzii* Fp360. Novel strain fp6 and *F. prausnitzii* Fp944 also formed a distinct cluster, which was significantly different from strain fp8 (bootstrap > 90). Strain fp5 formed a cluster with *F. prausnitzii* 28 and *F. prausnitzii* APC918_95b.

It is evident that novel strains fp1, fp2, fp3, fp7 and fp 9 are significantly different from the others and did not form close clusters with any of the novel strains. Furthermore, strains fp2, fp3, fp7 and fp9 did not cluster with any of the NCBI genomes. Although strain fp1 formed a cluster with NCBI genomes, it is notably distinct from them, as indicated by high bootstrap values.

### 3.4. Analysis of pharmabiotic potential

#### 3.4.1. Metabolomic analysis of SсFAs produced by novel strains of *F.prausnitzii*

A comparative analysis of the concentrations of ScFAs in the novel strains *F.prausnitzii* culture fluid was performed for 8 fatty acids: formic (FA), acetic (AA), propionic (PA), butyric (BA), isobutyric (iBA), 2-methyl-butyric (2mBA), valeric (VA), isovaleric (iVA). The structure of the analyzed data can be seen in Table 5. The average value (mean), standard deviation (std), minimum (min) and maximum (max) values, quartiles 1st and 3rd (Q1, Q3) are given.

**Table 5.** Data on concentrations of ScFAs produced by novel strains *F.prausnitzii* relative to an empty medium (control)

|  | AA | FA | PA | iBA | BA | iVA | 2mBA | VA |
| --- | --- | --- | --- | --- | --- | --- | --- | --- |
| **mean** | 0,73 | 2,18 | 1,39 | 0,67 | 16,27 | 0,82 | 0,81 | 2,76 |
| **std** | 0,48 | 1,68 | 1,70 | 0,14 | 3,74 | 0,08 | 0,07 | 3,28 |
| **Q1** | 0,34 | 1,58 | 0,79 | 0,56 | 13,12 | 0,76 | 0,75 | 1,29 |
| **Q3** | 0,96 | 1,88 | 0,94 | 0,75 | 19,76 | 0,87 | 0,86 | 2,22 |
| **min** | 0,15 | 1,08 | 0,73 | 0,50 | 12,30 | 0,72 | 0,72 | 1,09 |
| **max** | 1,74 | 6,89 | 6,23 | 0,87 | 22,11 | 0,93 | 0,92 | 11,98 |

*F. prausnitzii* actively produces butyrate (Equations 1-7), primarily through the consumption of external acetate. The synthesis of SсFAs requires the substrate Acetyl-CoA. The cell can obtain it from the breakdown of pyruvate (Equation 1) or from external acetate (Equation 2.2). In the process of pyruvate breakdown, formate is produced, which is a side product of butyrate biosynthesis. Furthermore, a portion of Acetyl-CoA can be converted to acetate (Equation 2.1), which is a significant product among SсFAs. However, this conversion can reduce the butyrate yield due to the additional consumption of Acetyl-CoA.

(1) Pyruvate + CoA-SH → Acetyl-CoA + **Formate**
(2.1) Acetyl-CoA → **Acetate** + ATP
(2.2) **Acetate** + CoA + ATP → Acetyl-CoA + AMP + Ppi
(3) 2 Acetyl-CoA ⇄ Acetoacetyl-CoA + CoA-SH
(4) Acetoacetyl-CoA + NADH + H⁺ → 3-Hydroxybutyryl-CoA + NAD⁺
(5) 3-Hydroxybutyryl-CoA → Crotonyl-CoA + H₂O
(6) Crotonyl-CoA + NADH → Butyryl-CoA + NAD⁺
(7) Butyryl-CoA + Pi + ADP → **Butyrate** + CoA + ATP

Additionally, *F. prausnitzii* can obtain propionate from oxaloacetate via transformations in the citrate cycle (Equations 8-14). Moreover, from Acetyl-CoA and Propionyl-CoA, the cell can produce valerate (Equations 15-16).

(8) Oxaloacetate + NADH + H⁺ → Malate + NAD⁺
(9) Malate → Fumarate + H₂O
(10) Fumarate + 2[H] → Succinate
(11) Succinate + CoA + ATP → Succinyl-CoA + ADP + Pi
(12) Succinyl-CoA → Methylmalonyl-CoA
(13) Methylmalonyl-CoA → Propionyl-CoA
(14) Propionyl-CoA + H₂O → **Propionate** + CoA
(15) Propionyl-CoA + Acetyl-CoA + 2 NADH + 2 H⁺ → Valeryl-CoA + CoA + 2 NAD⁺
(16) Valeryl-CoA + H₂O → **Valerate** + CoA

Thus, during the synthesis of butyrate, we can obtain formate and acetate as side products. Moreover, the more *F. prausnitzii* strives to obtain acetyl-CoA for the synthesis of butyrate, the more it will receive formate as a side product and the more it will consume acetate from external sources.

Hierarchical clustering based on ScFA production profiles revealed distinct metabolic groupings among the isolates. Strains fp2 and fp6 exhibited highly similar ScFA profiles, closely related to fp3, which showed the highest acetate concentration (6190 μM, 1.7-fold higher than control). Strains fp3 and fp4 shared comparable patterns of branched-chain acids (iBA, iVA, 2mBA), whereas fp4 displayed markedly elevated levels of propionate (1333 μM) and valerate (85 μM), exceeding control values by 6- to 12-fold. Strains fp5, fp7, and fp9 clustered together and were characterized by enhanced butyrate production, with fp9 identified as the most active producer. In contrast, fp1 and fp10 formed separate branches: fp10 exhibited generally low metabolite levels, whereas fp1 produced exceptionally high formate (9588 μM, sevenfold above control) along with increased acetate and butyrate synthesis. (Figure 7).

**Figure 7.**
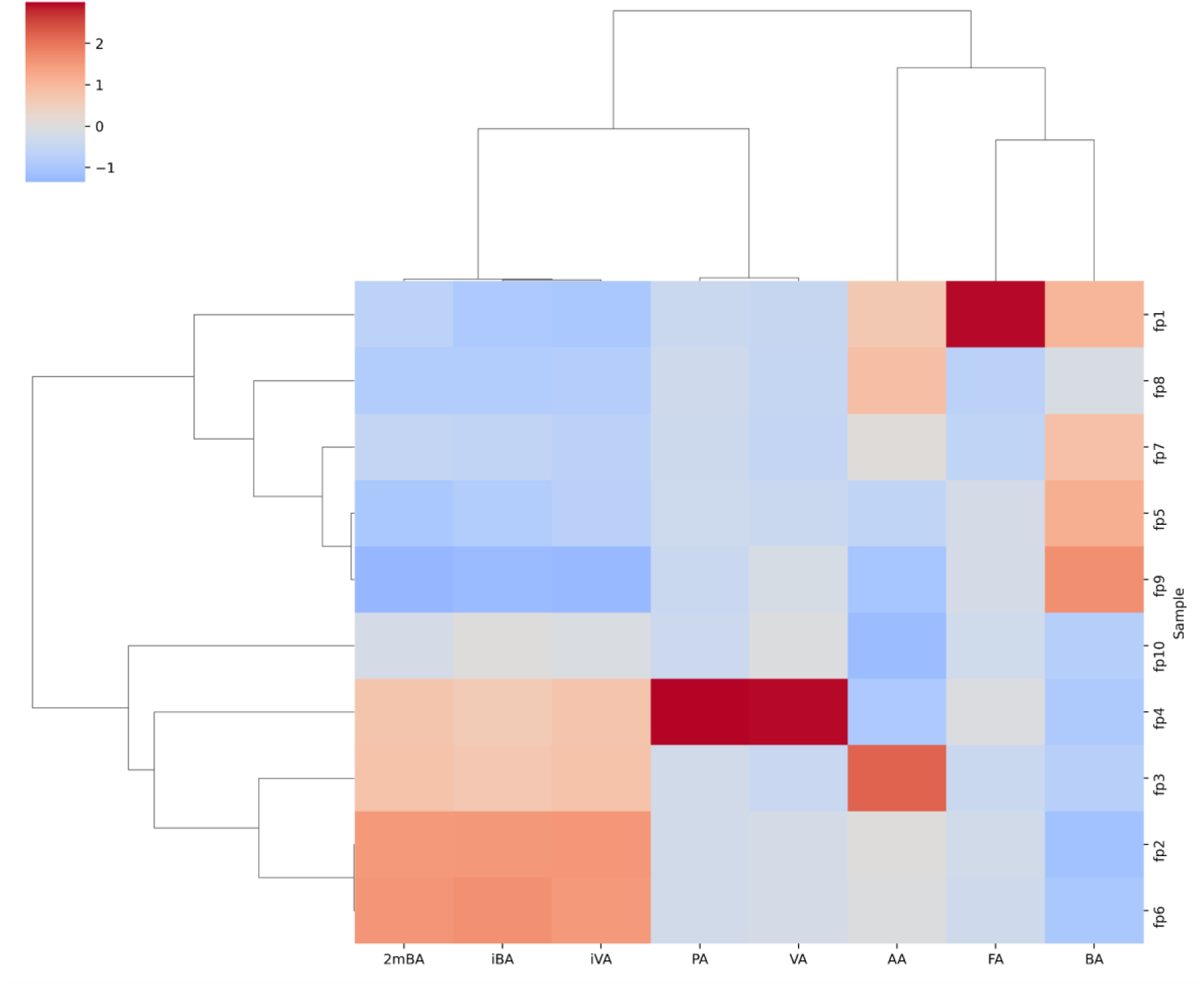
Clustermap of novel strain *F.prausnitzii* with hierarchic clustering of strains by metabolic profile, as well as with clustering of the acid profile by strain. Red (> 0), means that the relative acid content is higher than the average for the sample; Blue (< 0), means that the relative content is lower than the average for the sample. Close to beige (∼0) means that the relative content corresponds to the average of the sample.

The barplot comparing the concentrations of butyrate, acetate, and formate clearly demonstrates that the strains produce these SсFAs at different rates (Fig. 8).

**Figure 8.**
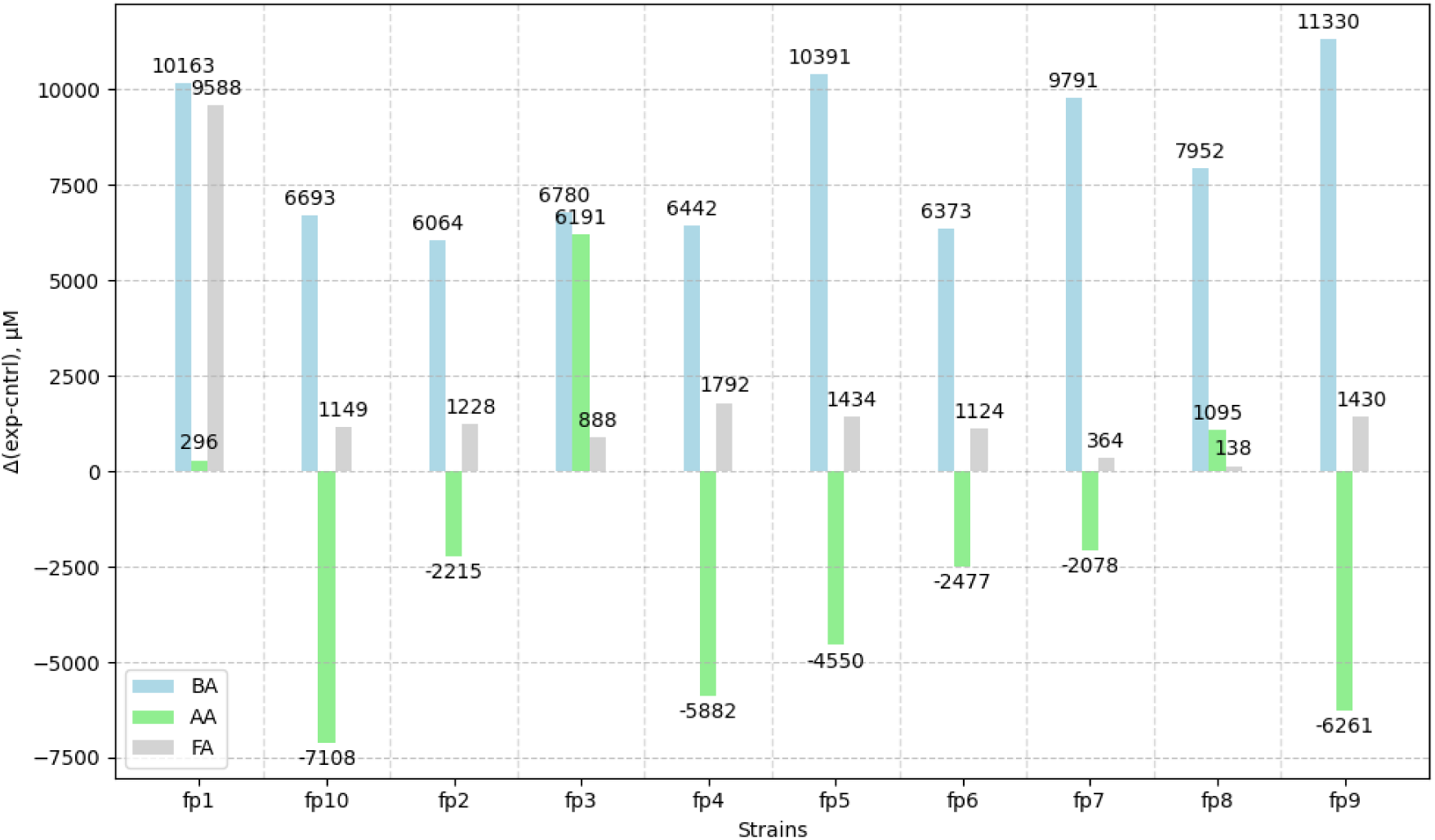
Barplot of butyrate (BA), acatate (AA), formate (FA) concentrations for novel strains *F.prausnitzii*. The X-axis shows 10 novel strains *F.prausnitzii*. The Y-axis shows the difference (µM) between the absolute ScFAы concentration values in the culture fluid of the strain and the values in the empty medium (control). The blue bars represent BA, the green bars represent AA, and the gray bars represent FA.

Strains fp9 and fp5, the highest butyrate producers, consumed ∼5000 µM acetate and produced >1000 µM formate, indicating preferential use of exogenous acetate for acetyl-CoA. Conversely, strain fp1 achieved similarly high butyrate yields with net acetate production and the highest formate levels, suggesting a primary reliance on pyruvate-derived acetyl-CoA.

Based on their distinct SсFA production profiles, strains fp1, fp4, and fp9 emerge as the most promising candidates for further study. Strain fp1 demonstrated high butyrate synthesis concomitant with significant formate and acetate production, indicating a primary reliance on pyruvate-derived acetyl-CoA. In contrast, strain fp4, while producing butyrate at a moderate level, uniquely generated notable quantities of propionate and valerate, SсFAs with recognized importance for gut health. Among the novel strains, fp9 was the most active dedicated butyrate producer, achieving the highest yield through the efficient utilization of extracellular acetate.

#### 3.4.2. Genomic analysis of novel strains according to reference genes catalogue

A comprehensive metabolite list was compiled (Supplementary Table 1). The catalogue of biologically active protein sequences, based on this list, annotates a total of 7,184 sequences of 264 enzyme genes derived from various genera described in our previous study. Homologs of these catalogue genes were identified across the genomes of all analyzed strains (Supplementary Table 2).

##### 3.4.2.1. Genes for ScFA metabolism

According to the results of a search for ScFA metabolism genes, 12 genes were found in the catalog (Table 6).

**Table 6.** Table describing the genes regulating the metabolism of ScFA found in novel strains.

| Gene | Reaction | Novel strains |
| --- | --- | --- |
| 3-hydroxyacyl-[acyl-carrier-protein] dehydratase (FabZ) | $(\beta\text{-hydroxyacyl-[acyl-carrier-protein]}) \rightarrow \text{trans-2-enoyl-[acyl-carrier-protein]} + \text{H}_2\text{O}.$ | All |
| 3-hydroxybutyryl-CoA dehydrogenase | $(\text{S})\text{-3-hydroxybutanoyl-CoA} + \text{NAD}^+ \rightleftharpoons \text{Acetoacetyl-CoA} + \text{NADH} + \text{H}^+.$ | All |
| Acetyl-CoA acetyltransferase (thiolase) | $2 \text{ Acetyl-CoA} \rightleftharpoons \text{Acetoacetyl-CoA} + \text{CoA-SH}.$ | All |
| Acyl-CoA dehydrogenase (short-chain specific) | $\text{Acyl-CoA} + \text{ETF(ox)} \rightarrow \text{Trans-2-enoyl-CoA} + \text{ETF(red)}.$ | All |
| Formate-tetrahydrofolate ligase | $\text{Formate} + \text{Tetrahydrofolate} + \text{ATP} \rightarrow 10\text{-formyl-THF} + \text{ADP} + \text{Pi}.$ | All |
| Methylmalonyl-CoA carboxyltransferase (5S, 12S) | $(\text{S})\text{-Methylmalonyl-CoA} + \text{Pyruvate} \rightleftharpoons \text{Propanoyl-CoA} + \text{Oxaloacetate}.$ | All |
| Butyryl-CoA dehydrogenase | $\text{Butyryl-CoA} + \text{ETF(ox)} \rightarrow \text{Crotonyl-CoA} + \text{ETF(red)}.$ | All |
| Butyryl-CoA:acetate CoA-transferase | $\text{Butyryl-CoA} + \text{Acetate} \rightarrow \text{Butyrate} + \text{Acetyl-CoA}$ | All |
| Phosphotransacetylase | $\text{Acetyl-CoA} + \text{Pi} \rightleftharpoons \text{Acetyl-phosphate} + \text{CoA-SH}.$ | All |
| Pyruvate:ferredoxin oxidoreductase | $\text{Pyruvate} + \text{CoA-SH} + 2 \text{ Fd(ox)} \rightarrow \text{Acetyl-CoA} + \text{CO}_2 + 2 \text{ Fd(red)} + \text{H}^+.$ | All |
| Pyruvate formate-lyase / Formate acetyltransferase | $\text{Pyruvate} + \text{CoA-SH} \rightarrow \text{Acetyl-CoA} + \text{Formate}.$ | fp1, fp2, fp3, fp6, fp8, fp9 |

The results clearly indicate the presence of butyrate, acetate, and formate metabolism in all examined strains. All strains possess the gene for phosphotransacetylase, which catalyzes a key step in the acetate production pathway from acetyl-CoA. They also encode formate-tetrahydrofolate ligase, responsible for consuming formate to generate 10-formyl-THF, thereby incorporating it into the biosynthesis of other compound groups.

Notably, the gene for pyruvate:ferredoxin oxidoreductase (PFOR), which generates acetyl-CoA from pyruvate via a redox reaction without producing formate as a byproduct, was detected in all strains. Conversely, the enzyme pyruvate formate-lyase (PFL), which catalyzes a similar reaction but yields formate, was not detected in strains fp4, fp5, fp7, and fp10. However, these strains still produced measurable quantities of formate. This suggests that strains fp4, fp5, fp7, and fp10 likely generate formate through an alternative pathway, such as via the breakdown of 10-formyl-THF. It is noteworthy that the pyruvate formate-lyase enzyme was also not detected in most NCBI strains (Figure S3).

We also performed a comparison of the homologs of the detected genes among the studied *F. prausnitzii* strains (Figure S1). This analysis revealed that strain fp1 exhibits relatively low homology (less than 97%) in the genes *fabZ, acetyl-CoA acetyltransferase, butyryl-CoA:acetate CoA-transferase* and *formate-tetrahydrofolate ligase* compared to the same genes in other strains. This suggests genetic distinctions in strain fp1 that may contribute to its high production of butyrate and formate. For these sequences in fp1, more than 10 amino acid substitutions were identified relative to the most common variant found in the other strains. These substitutions are single-point mutations located outside of known active sites. An insertion Gln-Gly-Lys was identified at position 54 in the *formate-tetrahydrofolate ligase* gene of strain fp1. Strain fp4 shows comparable differences in this gene group. These are likewise not associated with active-site regions and are mainly single-point mutations.

##### 3.4.2.2. Genes for gut homeostasis

Additionally, we performed an analysis of homologs involved in the metabolism of amino acids, polyamines, and anti-inflammatory substances (Table 7).

**Table 7.**
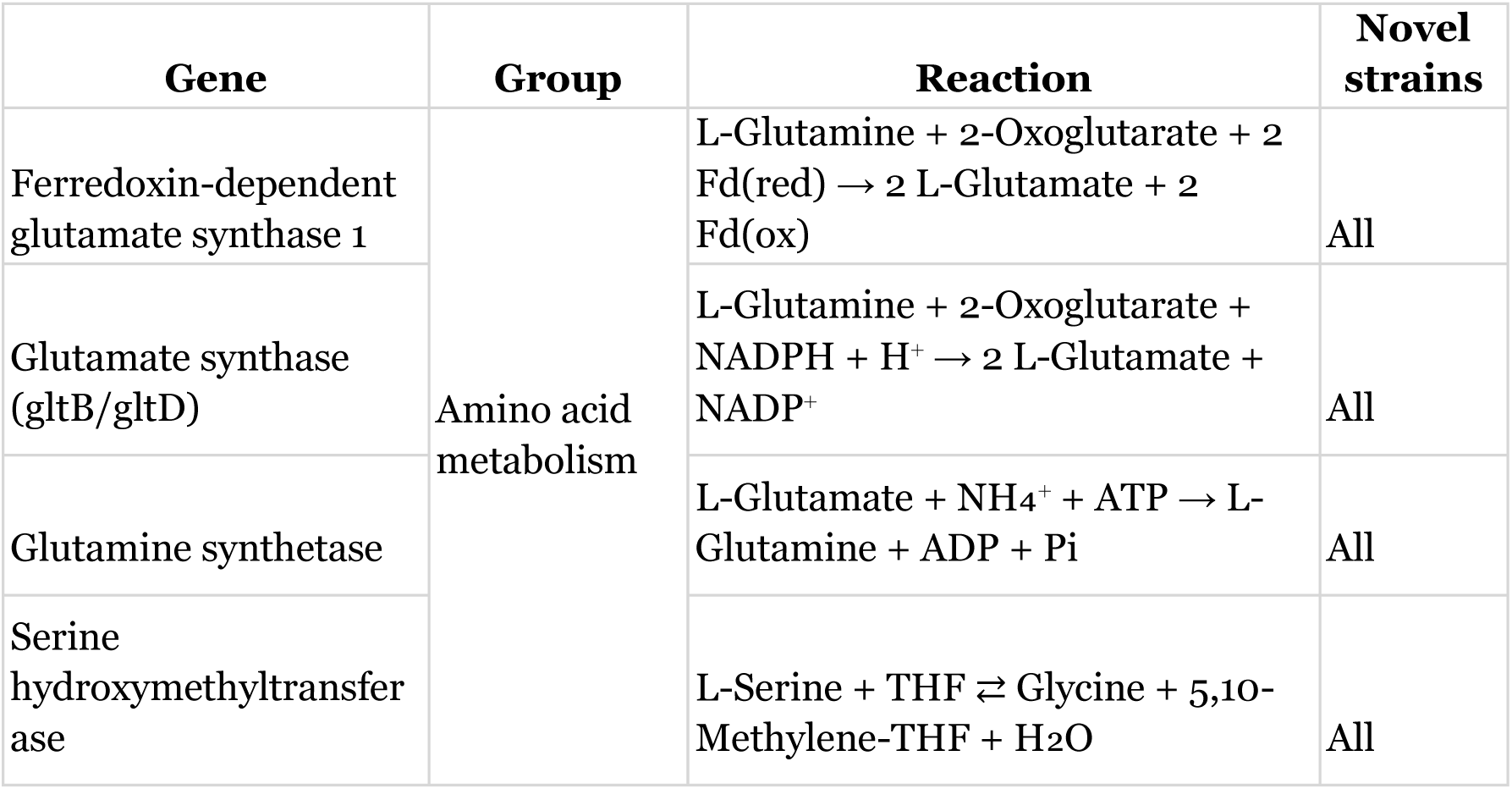
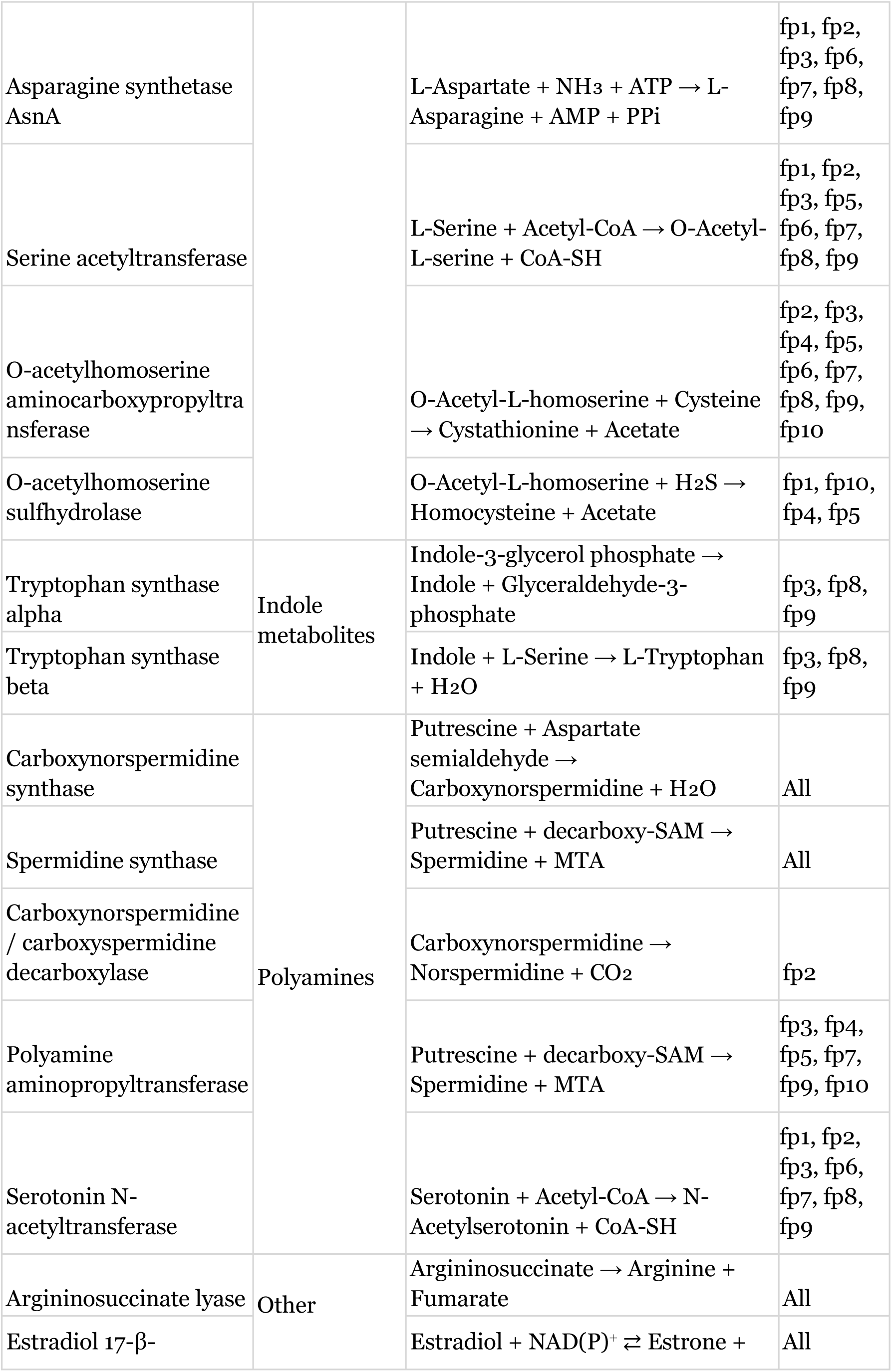
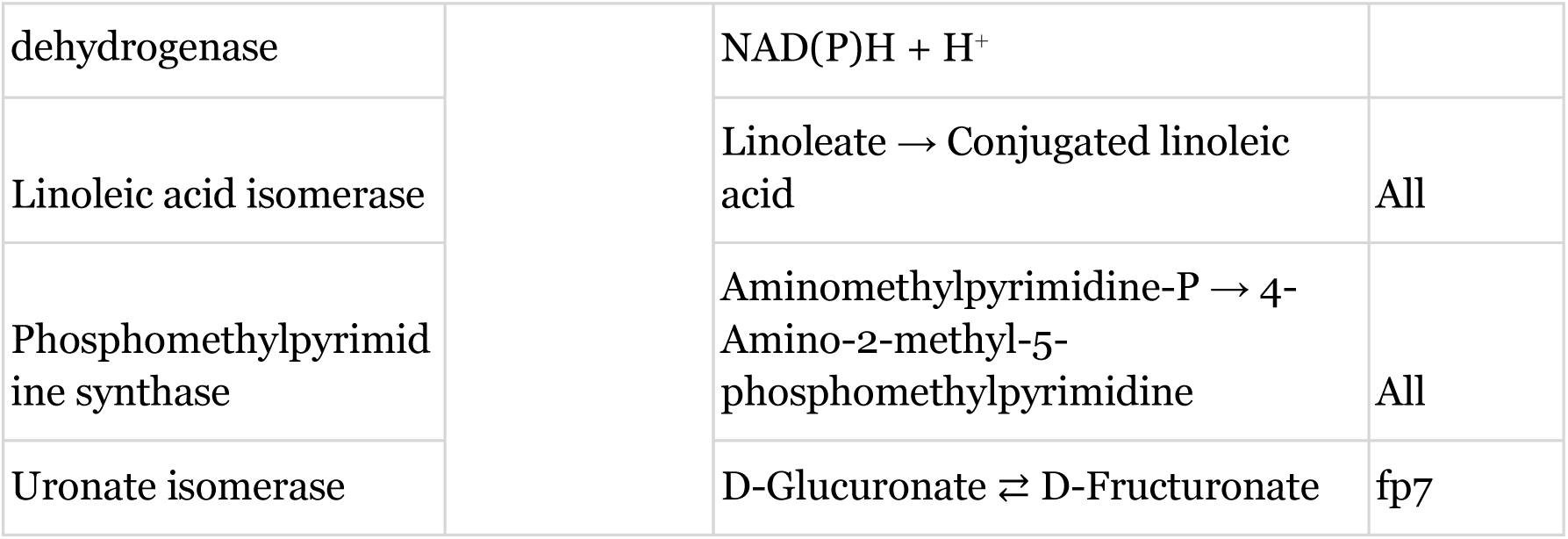
Table describing the genes regulating the metabolism of amino acids and other genes found in novel strains.

Notably, the repertoire of these genes differs among the strains fp1, fp4, and fp9, which were previously identified as the most promising candidates for pharmabiotics. Specifically, genes for *asnA, serine acetyltransferase*, and *serotonin N-acetyltransferase* were absent in strain fp4 but present in strains fp1 and fp9. Within this leading group, the *tryptophan synthase alpha/beta* genes were unique to strain fp9, while the microbial anti-inflammatory molecule gene was found exclusively in strain fp1. At the same time, the genes encoding *tryptophan synthase alpha/beta* were not detected for NCBI strains, and the microbial anti-inflammatory molecule gene was detected in only 4 NCBI strains (Figure S3). Overall, strains fp1 and fp9 possess the highest total number of genes regulating the metabolism of these important compounds. It is noteworthy that for most of the genes discussed in this section, the homology between some strains is low. In some cases, it is at the level of 80% (Figure S2).

##### 3.4.2.3. Genes of antibiotic resistance and virulence

A homology search against the Virulence Factor Database (VFDB) and the Comprehensive Antibiotic Resistance Database (CARD) identified 14 virulence-associated genes and one antibiotic resistance gene across the studied strains (Table 8).

**Table 8.** Table describing the genes from VFBD and CARD found in novel strains.

| Gene | Function | Novel strains |
| --- | --- | --- |
| clpP | Stress response, virulence regulation | All |
| groEL | Immunogenicity, stress tolerance | All |
| tufA | Adhesion, interaction with the immune system | All |
| galE | Capsule and LPS synthesis | All |
| wbtL | CPS precursors | All |
| cap8E | Capsular polysaccharide biosynthesis | fp1, fp2, fp4, fp6 |
| cps4K | Capsular polysaccharide biosynthesis | fp1, fp2, fp4, fp6 |
| cps4L | Capsular polysaccharide biosynthesis | fp1, fp2, fp4, fp6 |
| cpsI | Capsular polysaccharide biosynthesis | fp9 |
| glf | Capsular polysaccharide biosynthesis | fp1, fp2, fp3, fp5, fp6, fp7, fp8, fp9 |
| GBS_RS06585 | CPS / LPS formation | fp1, fp2, fp3, fp6 |
| wbtB | CPS synthesis | fp2, fp3, fp4, fp6, fp8, fp9 |
| ugd | Capsular polysaccharide biosynthesis | fp1, fp2, fp4, fp7 |
| hasB | Hyaluronic capsule biosynthesis | fp8 |
| tet(W) | Tetracycline resistance | fp1, fp2, fp4, fp6, fp9 |

It is noteworthy that genes such as *clpP, groEL*, and *tufA*, while included in VFDB, encode essential housekeeping functions (protease subunit, chaperonin, and elongation factor, respectively) and are not direct virulence factors. Their presence likely contributes to general environmental stress resistance. Another group of genes (*galE, wbtL, glf, wbtB, ugd, hasB*) is involved in utilizing host dietary carbohydrates and constructing the bacterial cell wall, facilitating gut colonization without inducing inflammation. Genes associated with capsule synthesis and phagocytosis avoidance (*cap8E, cps4K, cps4L, cps4I*) were found in several strains. Notably, only one of these genes was present in strain fp9. The tetracycline resistance gene *tet(W)* was detected in multiple strains, including fp1, fp4, and fp9, enhancing their survival potential in the presence of tetracycline antibiotics. For context, *F. prausnitzii* genomes from NCBI were found to harbor additional capsule genes (*cps4E, cps4I*) and more antibiotic resistance genes (against tetracycline, chloramphenicol, and macrolides).

## Discussion

This study investigates the genetic diversity and compares the pharmacobiotic potential of *Faecalibacterium prausnitzii* strains isolated from infants under one year of age in the Russian Federation and from adults worldwide. The research is based on the genetic characterization of all strains, supplemented by metabolomic analysis of strains from Russian infants. Our work presents, for the first time, data on *F. prausnitzii* from infant samples, whereas previously only genomic data from strains isolated from adults were available in the scientific literature. Furthermore, as part of this research direction, the first specialized solid medium for cultivating *F. prausnitzii* and other anaerobic organisms was developed. The medium is patented under Russian Federation Patent for Invention No. 2833071, issued on 14.01.2025; priority date 24.05.2024.

It has been found that, unlike bifidobacteria and lactobacilli, whose dominant species in the gut microbiota change with human age [57,58], *F. prausnitzii* is detected in samples from both infants and adults. It is known that prior colonization by other microbes is necessary for the reliable detection of *F. prausnitzii* in early life [59]. It has also been suggested that gut colonization by *F. prausnitzii* requires an oxygen-deficient gut environment or, presumably, another type of environmental exposure mediated by other microbes [60].

Sequencing of *F. prausnitzii* is also a crucial task for studying its genomes and properties [61]. In our study, two types of reads were generated: long reads obtained using the Oxford Nanopore MinION platform and short reads obtained using the Illumina HiSeq 2500 platform. Through the combined use of these reads and *de novo* genome assembly methods, we obtained 4 complete and 6 draft genomes of *F. prausnitzii*. Obtaining complete genomes for this species is a non-standard and challenging task [62].

A comprehensive characterization of the novel *F. prausnitzii* genomes was performed. To confirm taxonomic assignment to the species *F. prausnitzii*, two independent methods were employed. The results of both methods were consistent, confirming that the studied strains belong to the species *F. prausnitzii*. However, when analyzing ANI metrics for the same strain, the highest similarity scores were recorded against different reference genomes. This indicates that each metric reflects different aspects of genomic similarity, ranging from shared nucleotide identity to structural and compositional features of the genome. The absence of a single reference genome with the highest ranking across all metrics underscores the intraspecific genomic heterogeneity of *F. prausnitzii*. It also highlights the necessity of a comprehensive approach using multiple metrics for robust taxonomic assignment and comparative genomic analysis.

Furthermore, the comparison of nucleotide sequences from the novel genomes yielded non-identical results between ANI-based metrics and core-gene phylogeny. Based on ANI metrics, the most similar strain pairs were fp1/fp2, fp4/fp10, fp3/fp7, fp3/fp9, fp7/fp9, and fp6/fp8 (Figure 2). In contrast, the core-gene phylogenetic analysis, which included reference strains from NCBI, formed clusters only for the pairs fp4/fp10 and fp6/fp8. Strain fp1 clustered exclusively with certain NCBI strains, while strains fp2, fp3, fp7, and fp9 did not form distinct clusters with each other or with other strains. Notably, the phylogenetic branches for strains fp3, fp7, and fp9 were positioned close to one another (Figure 6). Analysis of genes against a reference catalog revealed that for most homologs, strains fp3, fp6, fp7, fp8, and fp9 exhibited the highest homology percentages. A similar pattern was observed for strains fp4 and fp10 (Figures S1, S2).

A pangenomic analysis of new strains and strains with complete genomes from NCBI was also performed. An open pangene was obtained. According to his results, all strains have a large number of accessory genes, and the number of unique genes varies among all strains. The fp2, fp3, and fp6 strains are characterized by the lowest values of unique genes (more than 100). Such an open pangenome pattern is typical for gut commensal bacteria that inhabit diverse ecological niches and interact dynamically with other microbiota and the host environment. This genomic flexibility may underlie the observed functional diversity among *F. prausnitzii* strains and their adaptation to different intestinal conditions [63]. These results demonstrate that essential metabolic functions are conserved in the core genome, while environmental interaction and adaptive functions are encoded within the accessory and unique genomic components, providing a genetic basis for the ecological versatility of *F. prausnitzii*.

Based on the ANI metric, the strains were categorized into three groups: high similarity, medium similarity, and low similarity. In contrast, the phylogenetic tree constructed from core genes did not allow for a clear separation into up to three distinct groups, as it yielded a more complex and individualized clustering pattern. However, homologs analysis revealed that strains fp3, fp6, fp7, fp8, and fp9 formed a distinct cluster. Strains fp4 and fp10 constituted another separate group. The remaining strains could be tentatively classified as a third group, although they did not exhibit high homology scores among themselves.

The metabolomic analysis of ScFAs also enabled strain grouping. The cluster map (Figure 7) shows that the most similar acid profiles are shared by strains fp2, fp6 and, separately, by strains fp5, fp7, fp9. Strains fp4 and fp10 exhibit a notably different ScFA quantity profile. However, when examining the absolute concentrations of formic, acetic, and butyric acids (Figure 8), the profiles of strains fp4 and fp10 appear almost identical. They are characterized by a medium level of butyric acid (concentrations around 6000 µM), a low level of formate (around 1000 µM), and the consumption of over 5000 µM of acetate. Overall, these ratios of formate, acetate, and butyrate characterize strains fp4 and fp10 as not the most active butyrate producers; despite high acetate uptake, they demonstrate only moderate butyrate levels. A similar pattern is observed for strains fp2 and fp6. However, they consumed less acetate (around 2000 µM) while maintaining butyrate and formate levels comparable to strains fp4 and fp10. This indicates a higher relative activity in butyrate synthesis. Strain fp8 also produced similar butyrate concentrations. Unlike the previous groups, it simultaneously synthesized acetate (approximately 1000 µM) and produced significantly less formate (only 100 µM). Acetate synthesis was also observed in strains fp1 and fp3. Strain fp1 is characterized by the highest formate concentration and one of the highest butyrate concentrations. Strain fp3 synthesized approximately equal amounts of butyrate and acetate (around 6000 µM each), along with about 1000 µM of formate. This suggests that fp3 follows a butyrate synthesis pathway similar to fp1, but at a slower rate. A similar assumption can be made for strain fp8. Strains fp5, fp7, and fp9 are characterized by high butyrate concentrations (over 10,000 µM) and high acetate consumption (around 5000 µM), unequivocally placing them in a single group. Among them, strain fp9 is the most active butyrate producer, yielding the highest butyrate concentration. Based on butyrate synthesis patterns, we can propose the existence of three functional groups: active butyrate producers with acetate consumption and negligible formate levels (fp5, fp7, fp9); less active butyrate producers (fp2, fp4, fp6, fp10); butyrate producers with concomitant acetate and formate synthesis (fp1, fp3, fp8), within this group, fp1 is the most active producer of both butyrate and formate. The division into 3 groups is also observed in other studies of *F.prausnitzii*, however, they also demonstrate significant heterogeneity, especially in the synthesis of butyrate, as in our case [64,65]

Furthermore, the novel investigated strains demonstrated a lower abundance of virulence and antimicrobial resistance genes compared to strains from the NCBI database. In addition to the genes listed in Table 8 that were identified in the new strains, the NCBI strains were found to possess genes such as *cps4E, cps4I,* as well as antibiotic resistance genes conferring resistance to tetracycline, chloramphenicol, and macrolides. Notably, the NCBI strains did not carry genes encoding the MAM or genes involved in tryptophan biosynthesis. We also identified several less common genes with specific metabolic functions. For instance, strain fp7 harbors a uronate isomerase, which may enhance its ability to metabolize dietary fibers in the host intestine. Strain fp2 encodes a carboxynorspermidine / carboxyspermidine decarboxylase, an enzyme involved in the synthesis of spermidine and norspermidine. In summary, the novel *F. prausnitzii* strains possess a reduced repertoire of putative virulence determinants and only one confirmed antibiotic resistance gene compared to reference strains. Concurrently, the studied strains are characterized by a higher prevalence of genes associated with the synthesis of metabolites linked to improved intestinal and host health (Figure S3). This genomic profile supports their potential suitability for development as pharmabiotics.

## Conclusions

Thus, we cultivated and isolated ten novel strains from infant feces, assembled their genomes, and performed comprehensive genomic characterization. Genomic and metabolomic profiling identified strains fp1 and fp9 as the most promising pharmabiotic candidates. Both are highly active butyrate producers, with strain fp1 additionally exhibiting high formate production. Furthermore, strain fp1 possesses the most extensive repertoire of beneficial genes while concurrently harboring the fewest virulence and pathogenicity determinants.

## Supporting information

https://drive.google.com/drive/folders/11pafgQGCt1YZyVmaJNj5QLf6iX3AYJv2?usp=drive_link

## Acknowledgements

The work was carried out with the support of the project of the Ministry of Health of Russia “Development of an integrated approach to the diagnosis and correction of dysbiotic disorders of the gut microbiota caused by antibacterial therapy in newborns” (124020600025-2).

## Author Contributions

Conceptualization: Galanova O.O., Vatlin A.A., Danilenko V.N.

Methodology: Kovtun A.S., Troshina D.A., Muravieva V.V.

Investigation: Galanova O.O., Akulinin M.D., Troshina D.A., Kovtun A.S., Odorskaya M.V., Ksenia N. Zhigalova, Roman V. Izyumov, Alexey B. Gordeev, Bayr O. Bembeeva, Elena L. Isaeva

Resources: Galanova O.O., Vatlin A.A.

Data curation: Galanova O.O.

Writing—original draft preparation: Galanova O.O., Troshina D.A.

Writing—review and editing: Galanova O.O., Kovtun A.S., Vatlin A.A., Bekker O.B.

Visualization: Galanova O.O., Akulinin M.D.

Supervision: Danilenko V.N., Tatiana V. Priputnevich, Gennady T. Sukhikh

Project administration: Vatlin A.A., Galanova O.O., Alexey B. Gordeev, Danilenko V.N.

Funding acquisition: Tatiana V. Priputnevich, Gennady T. Sukhikh

