## Supplementary figures and images for "Identification and Characterization of Novel *Faecalibacterium prausnitzii* Strains with Potential Pharmabiotic Applications"

### Figure S1.png

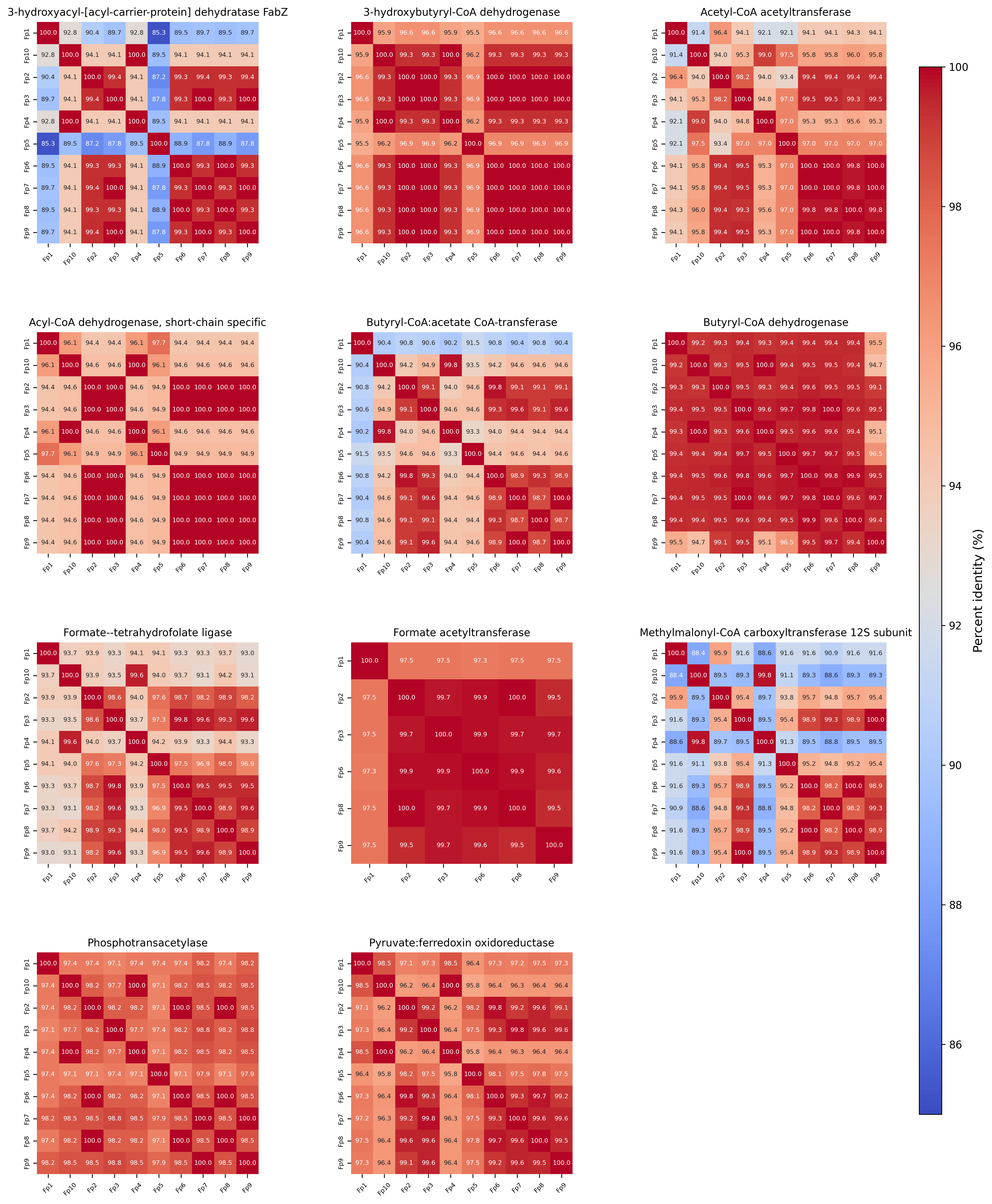

### Figure S3.png

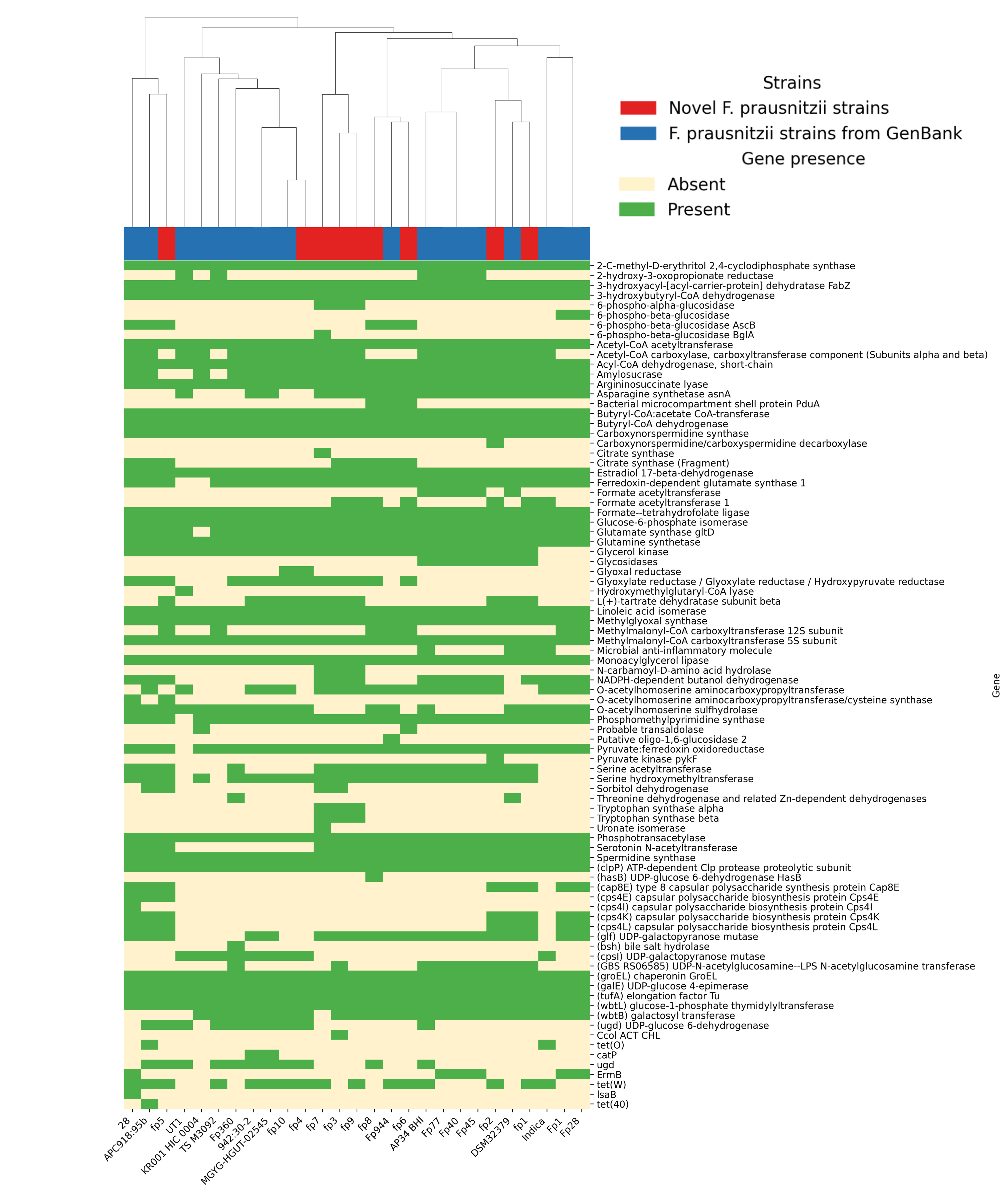
